# Characterization of the continuous transcriptional heterogeneity in Wilms’ tumors using unsupervised machine learning

**DOI:** 10.1101/2022.06.06.494924

**Authors:** Yaron Trink, Achia Urbach, Benjamin Dekel, Peter Hohenstein, Jacob Goldberger, Tomer Kalisky

## Abstract

Wilms’ tumors are pediatric malignancies that are thought to arise from faulty kidney development. To date, the course of treatment depends on manual histopathological classification which is difficult since the tumors differ between patients in a continuous manner. Here, we used three computational approaches to characterize the continuous heterogeneity of Wilms’ tumors. We first chose a published dataset of microarray gene expression measurements from high-risk blastemal-type Wilms’ tumors. Then, we used Pareto Task Inference to show that the tumors form a triangle-shaped continuum in latent space that is bounded by three tumor archetypes with “stromal”, “epithelial”, and “blastemal” characteristics, that resemble the un-induced mesenchyme, the Cap mesenchyme, and early epithelial structures of the fetal kidney. We confirmed this by fitting a generative probabilistic “grade of membership” model whereby each tumor is represented as a unique mixture of three hidden “topics” with blastemal, stromal, and epithelial characteristics. Finally, we used cellular deconvolution to show that each tumor is composed of a unique mixture of cell populations resembling the un-induced mesenchyme, the cap mesenchyme, and the early epithelial structures of the fetal kidney. We anticipate that these methodologies will pave the way for more quantitative strategies for tumor stratification and classification.

## INTRODUCTION

Wilms’ tumors, also known as nephroblastomas, are pediatric malignancies that develop in children under age five. They contain renal progenitor cells whose differentiation is incomplete and are therefore thought to be the result of faulty kidney development [1].

There are currently two main protocols for treating Wilms’ tumor patients. The protocol of the Children’s Oncology Group (COG), followed in North America, dictates primary surgery followed by chemotherapy [2]. The protocol of the Société International d’Oncologie Pédiatrique (SIOP) [3], followed in Europe and other countries, dictates preoperative chemotherapy followed by surgery, postoperative chemotherapy and radiotherapy. Although treatment outcomes are usually favorable, patients with bilateral, relapsed, and high-risk tumors are more difficult to treat. Moreover, survivors of childhood cancers are at increased risk for health problems later on in life [4]. Therefore, it is important to discern between low risk patients which have high survival rates and could benefit from a reduction in therapy, and the high-risk patients that require more aggressive treatment [5,6].

Since Wilms’ tumors arise from genetic and epigenetic distortions during different stages of fetal kidney development, they are highly heterogeneous. There are generally three “archetypical” forms of Wilms tumors: Stromal tumors contain a large fraction of cells that resemble the un-induced mesenchyme of the developing fetal kidney, and sometimes additional tissues such as muscle and cartilage that, like the un-induced mesenchyme, also originate from the fetal mesoderm [1]. Blastemal tumors resemble the cap mesenchyme, which is a transient cellular compartment in the fetal developing kidney that contains the nephron progenitors. Epithelial tumors contain a significant portion of early non-functional epithelial structures. However, in many cases the tumors contain a mixture of all three components and are classified as “mixed” or “triphasic”. The component proportions can vary significantly resulting in a continuous range of tumor appearances [7].

Histopathological classification is currently the gold standard for diagnosis and subtyping in both COG and SIOP protocols. For example, tumors with remaining viable blastema after being treated with preoperative chemotherapy according the SIOP protocol have worse prognosis [8]. However, precise classification can often be challenging for several reasons. First, many pathologists lack experience in this domain due to the rarity of cases [9]. Second, the similarity to other renal tumors often confounds proper classification [7]. Third, the varying morphology between cases makes the pathological diagnosis difficult to characterize and interpret since it is continuous in nature and therefore unsuited for compartmentalization into one of several discrete subtypes, especially in triphasic tumors.

Therefore, in this study we set to characterize the heterogeneity of Wilms’ tumors in a more quantitative and interpretable way using gene expression measurements and computational algorithms. We first chose a set of published microarray gene expression measurements of high-risk, blastemal type, post-operative chemotherapy Wilms’ tumors [8] and examined the geometry of their formation in latent space. We found that, rather than being arranged in discrete and well separated clusters, the tumors form a triangle shaped continuum in the space spanned by the first two principal components [10] (Figure 1). Using Pareto task inference [11] and GO enrichment analysis, we showed that the vertices of this triangle represent stromal, blastemal, and epithelial cellular “archetypes”, that correspond to the three main lineages in the developing kidney – the un-induced mesenchyme, the Cap mesenchyme, and the renal tubular epithelium (Figure 2). Next, in order to provide a more quantitative interpretation to this geometrical description of Wilms’ tumor heterogeneity, we used Topic modeling [12,13] to fit a generative probabilistic model to our dataset and showed that each tumor can be represented as a unique mixture of three hidden “topics” with blastemal, stromal, and epithelial characteristics (Figure 3). Finally, since the histology and cellular composition of Wilms’ tumors resembles that of the fetal kidney, we complemented the above analysis by performing cellular deconvolution [14] with respect to an independently measured single-cell gene expression dataset from a fetal kidney in order to model the bulk tumors in terms of their underlying single cell composition (Figure 4). Cellular deconvolution showed that each tumor can be represented as a unique mixture of cell populations resembling the un-induced mesenchyme, the Cap mesenchyme, and the early epithelial structures of the fetal kidney, thus inferring a detailed single-cell based profile for each tumor.

**Figure 1:**
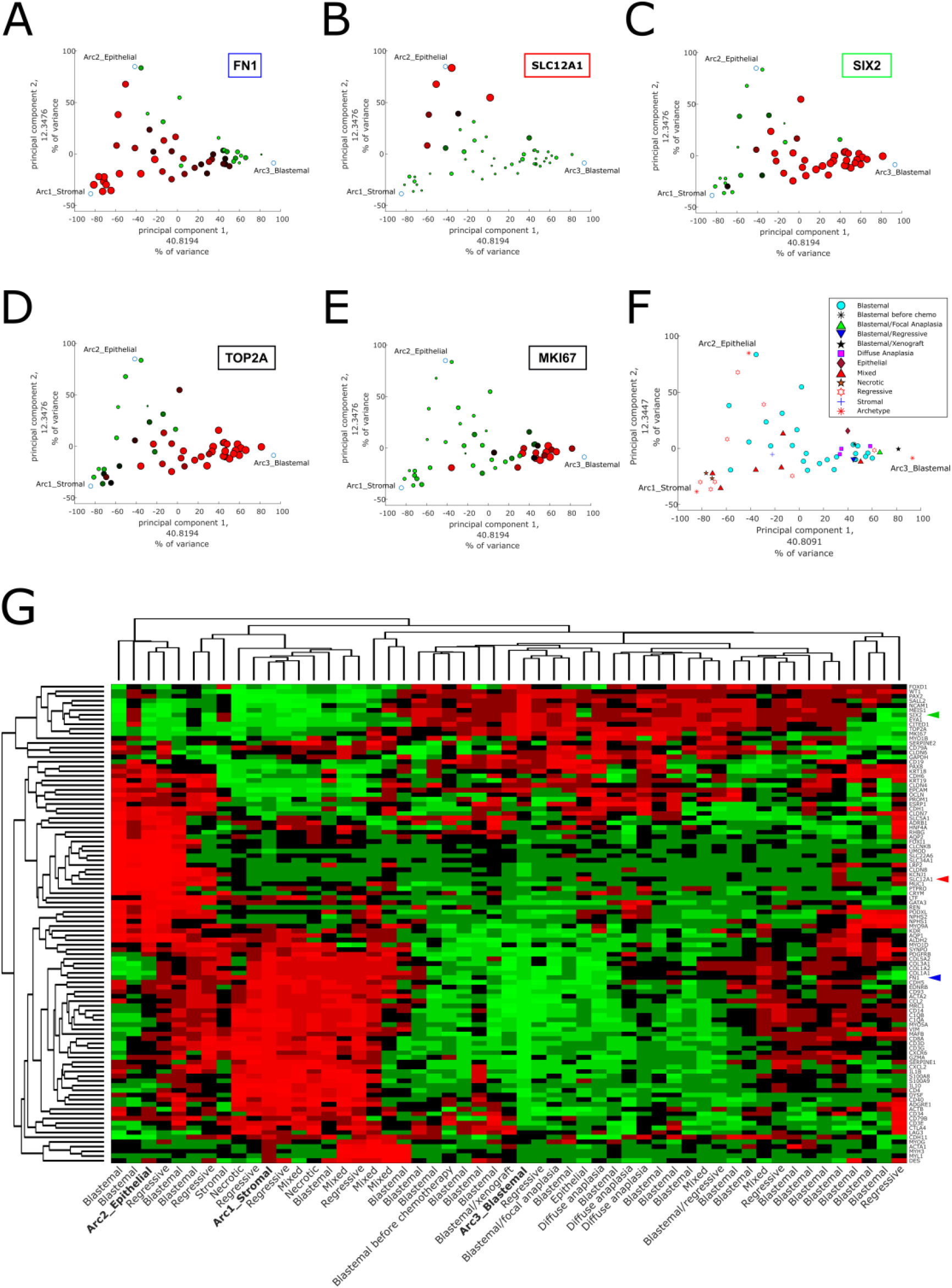
Pareto Task Inference shows that blastemal type, post-operative chemotherapy Wilms’ tumors fill a triangle shaped continuum in latent space that is bounded by archetypes with stromal, epithelial, and blastemal characteristics (A-E) Shown are PCA plots of the tumors in our dataset along with the three calculated archetypes (the vertexes of the triangle). In order to identify the archetypes, in each panel we selected a gene marking one of the three main lineages of the fetal developing kidney and plotted the size and color of each point according to its expression level. It can be seen than FN1, a marker for the un-induced mesenchyme, is highly expressed towards the stromal archetype. Likewise, SLC12A1, which marks the renal epithelium, is highly expressed near the epithelial archetype, and SIX2, a marker for the Cap mesenchyme, is predominantly expressed near the blastemal archetype. The genes TOP2A and MKI67, which mark cycling cells, are also highly expressed near the blastemal archetype. (F) A PCA plot of the tumors marked according to their reported histological type. It can be seen that tumors with anaplastic histology (diffuse or focal), which is considered least favorable, tend to cluster in the vicinity of the blastemal archetype. (G) A gene expression heatmap of 102 genes that are known from the literature to mark specific kidney lineages. The three arrowheads mark the genes from panels A, B, and C. It can be seen that the stromal, blastemal, and epithelial archetypes over-express genes that mark the un-induced mesenchyme, the Cap mesenchyme, or renal epithelial tubular structures, respectively. Hierarchical clustering was done using the MATLAB clustergram function with standardized rows (=genes), Euclidean distance metric, and average linkage.

**Figure 2:**
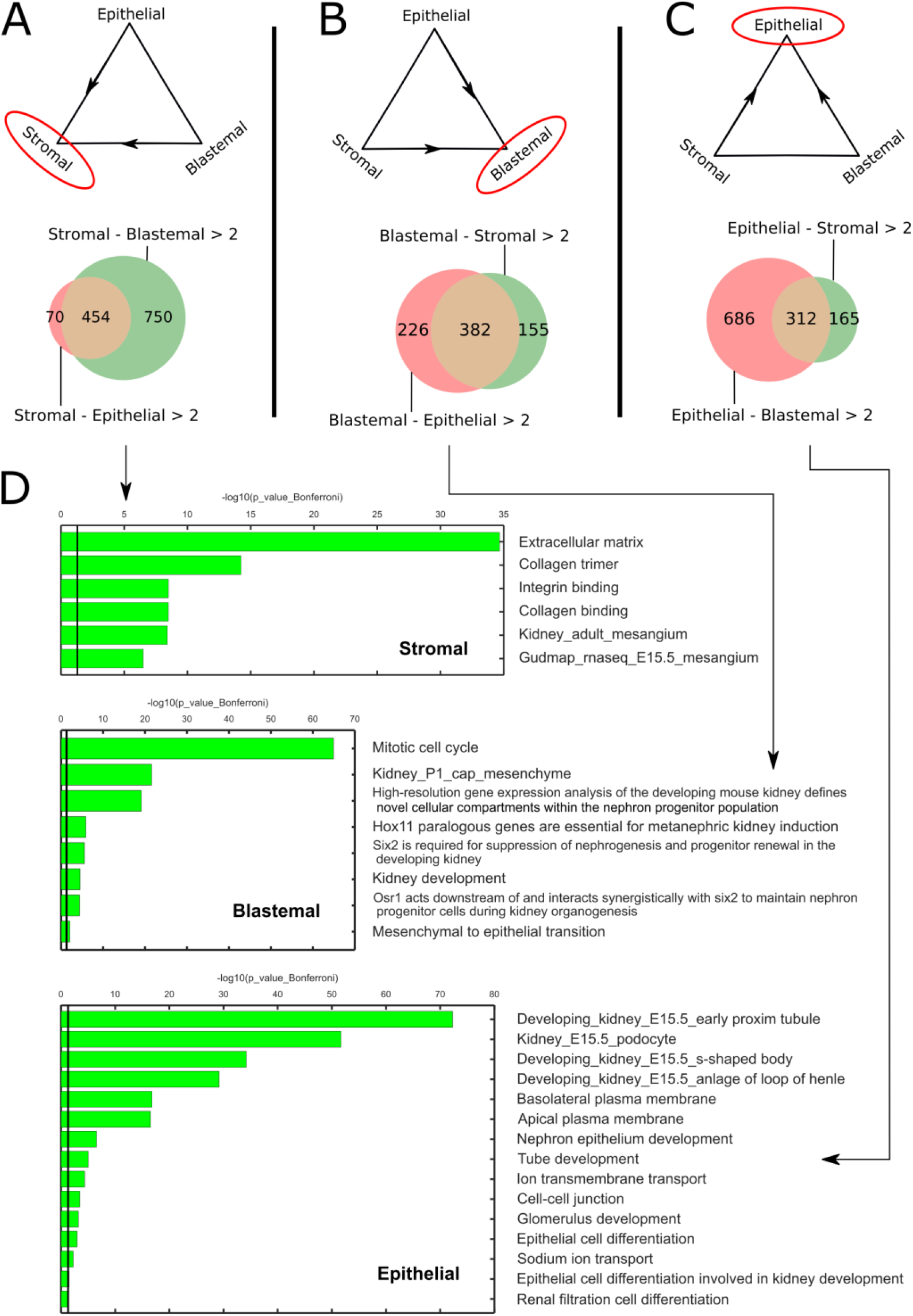
GO enrichment analysis shows that the stromal, blastemal, and epithelial archetypes are enriched for genes that are characteristic of the un-induced mesenchyme, the Cap mesenchyme, and renal epithelium of the fetal developing kidney, respectively. (A-C) For each of the three archetypes, we selected a set of genes for which the log2-fold change was larger than two with respect to the other two archetypes. These genes were used as input to Toppgene. (D) It can be seen that the stromal archetype over-expresses genes that are characteristic of the un-induced mesenchyme, for example, genes responsible for creating and maintaining the extracellular matrix. The blastemal archetype over-expresses genes that are characteristic of the Cap-mesenchyme, for example, genes responsible for maintaining nephron progenitors and for the mesenchymal to epithelial transition (MET). Likewise, the epithelial archetype over-expresses genes that are characteristic of the fetal renal epithelium, for example genes responsible for nephron epithelial differentiation and genes responsible for creation and maintenance of renal proximal tubules, S-shaped bodies, cell-cell junctions, and membrane transporters.

**Figure 3:**
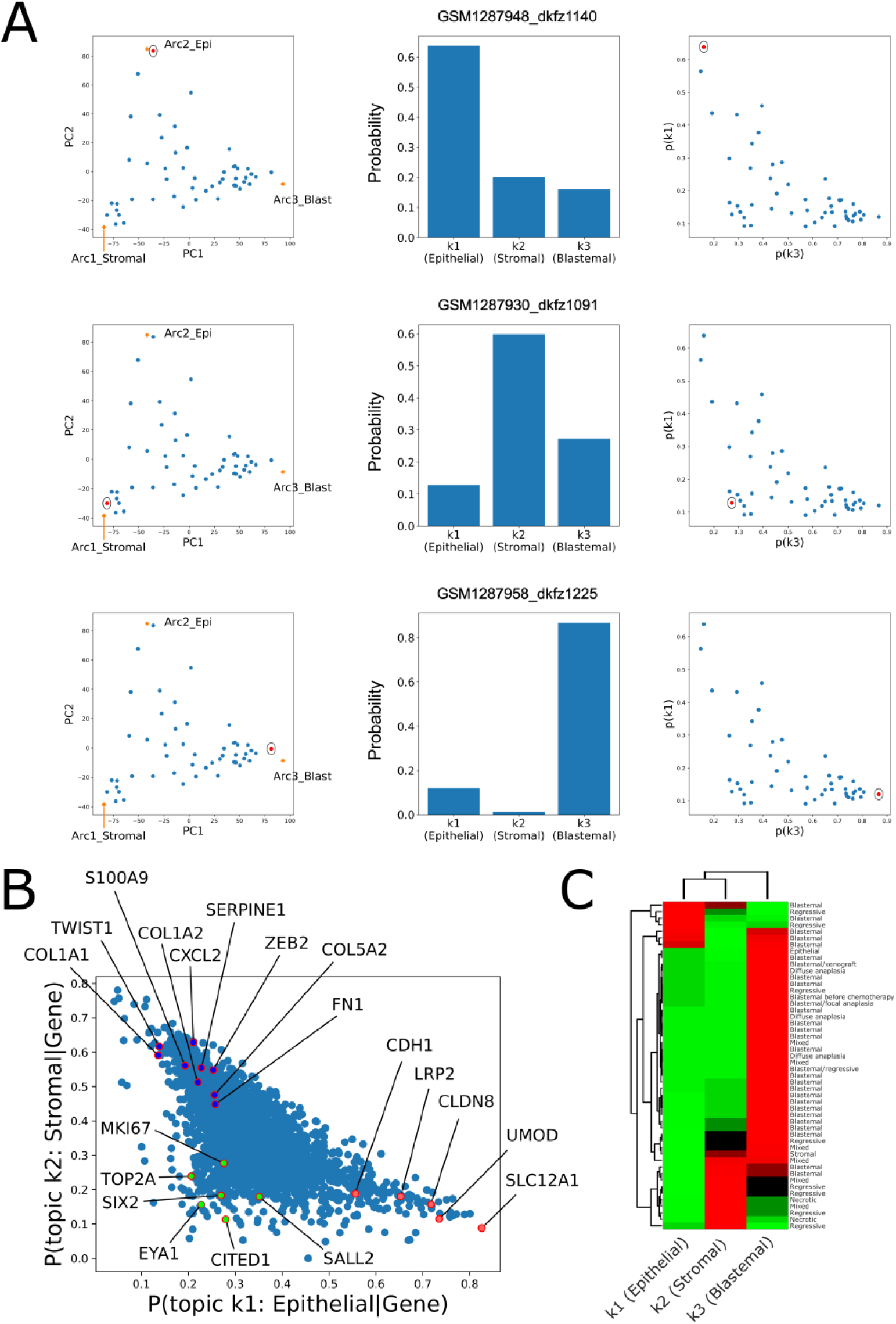
Topic modeling shows that each tumor can be represented as a unique mixture of three hidden topics with blastemal, stromal, and epithelial characteristics. (A) Shown are the fractions of topics (center and right panels) for selected tumors that are located near each of the three archetypes in latent space (left panels). The topic model predicts that each one of these tumors was generated primarily from a single topic. In particular, tumors near the epithelial topic contain a high fraction of topic no. 1 (top panels), tumors near the stromal topic contain a high fraction of topic no. 2 (middle panels), and tumors near the blastemal topic contain a high fraction of topic no. 3 (bottom panels). It can also be seen that the geometry of the topic simplex (right panels) resembles that of the PCA latent space (left panels). (B) The posterior probabilities of the three topics given the expression of specific genes show that the three topics have stromal, blastemal, and epithelial characteristics. The posterior probabilities of a topic given the expression of a specific gene represent the association between over-expression of that specific gene and the probability of that topic being represented in a tumor. It can be seen that that that over-expression of markers for the renal epithelial tubules (e.g. CDH1, SLC12A1, LRP2, and UMOD) is associated with high probability for the epithelial topic (topic no. 1), over-expression of markers for the un-induced mesenchyme (COL1A1, COL1A2, COL5A2, FN1, and SERPINE1) is associated with high probability for the stromal topic (topic no. 2), and over-expression of markers for the Cap mesenchyme (SIX2, CITED1, EYA1, and SALL2) is associated with high probability for the blastemal topic (topic no. 3). (C) A heatmap of the topic composition of each tumor. It can be seen that, tumors with anaplastic histology (diffuse or focal), which is considered least favorable, as well as the single blastemal xenograft in the dataset, all contain a large fraction of the blastemal topic.

**Figure 4:**
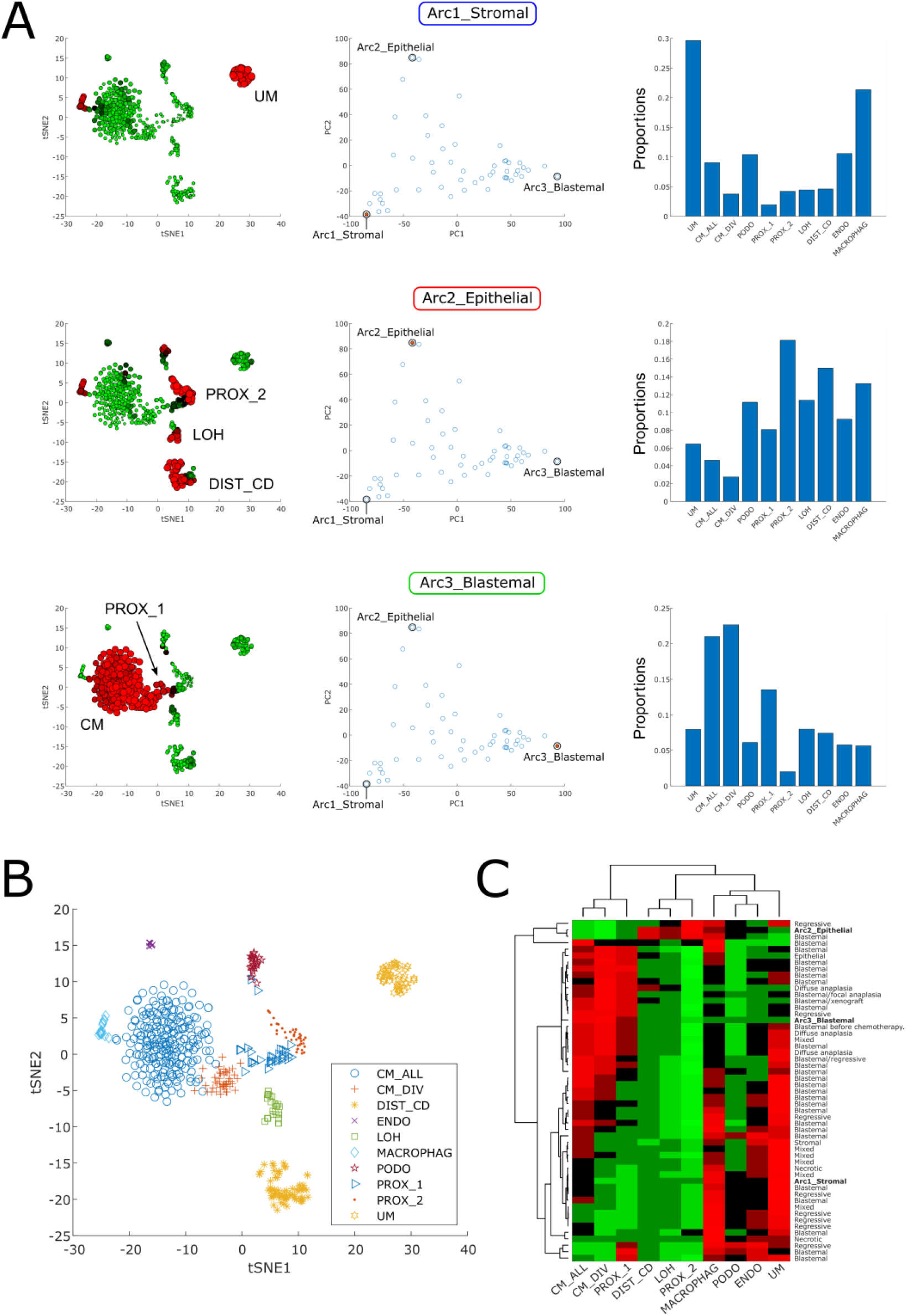
Cellular deconvolution indicates that each tumor is composed of a unique mixture of cell populations resembling the un-induced mesenchyme, the cap mesenchyme, or the early epithelial structures of the fetal kidney. (A) Shown is a cellular deconvolution of the stromal, blastemal, and epithelial archetypes. The left panels show tSNE plots of the reference single cell RNAseq dataset from the developing mouse fetal kidney that was used for the deconvolution. The size of each point (=single cell) corresponds to the proportion of that cell in the specific de-convolved tumor. The middle panels highlight the relevant tumor or archetype in the PCA latent space as a red dot. The right panels show barplots of the predicted proportions of each of the 10 cell types in the reference single-cell matrix. It can be seen that the stromal archetype is composed primarily from cells resembling the un-induced mesenchyme (UM) and macrophages, the epithelial archetype is composed mainly of epithelial tubular cells resembling those of the fetal proximal tubule (PROX_2), the Loop of Henle (LOH), and the distal tubule (DIST_CD), and the blastemal archetype is composed primarily from cells resembling the Cap Mesenchyme (CM), along with some very early epithelial tubular structures (PROX_1). (B) The different cell populations marked on a tSNE plot of the reference single cell RNAseq dataset from the developing mouse fetal kidney that was used for cellular deconvolution (CM – Cap mesenchyme, DIST_CD – distal tubule and collecting duct, ENDO – endothelial, LOH – Loop of Henle, MACROPHAG – macrophages, PODO – podocytes, PROX_1 – early epithelial structures such as C/S-shaped bodies, PROX_2 – proximal tubule, UM – un-induced mesenchyme). (C) A heatmap of the cell type proportions from which each tumor is composed, as predicted by cellular deconvolution. It can be seen that most of the tumors with reported blastemal histology contain a significant proportion of cells resembling those of the Cap mesenchyme (CM), as expected. Likewise, tumors with anaplastic histology (diffuse or focal), as well as the single blastemal xenograft in our dataset, are also composed mainly of cells resembling the cycling Cap mesenchyme cells (CM_DIV) in the fetal kidney. On the other hand, tumors with triphasic/mixed or regressive histology contain high proportions of cells resembling the un-induced mesenchyme (UM).

We anticipate that these results will lead to more quantitative and precise strategies for Wilms’ tumor classification and stratification that will eventually enable more personalized therapies for pediatric kidney malignancies.

## METHODS

### Gene expression datasets

53 CEL files were downloaded from the GEO database (accession number GSE53224). We also received a table connecting the microarray ID’s from the GEO database (GSM1287918_dkfz1079, GSM1287919_dkfz1080, …) with the tumor identifiers from Table S2 in the original publication by [8] (WT055, WT056, …) from Prof. Manfred Gessler, who is one of the authors of the original study (Table S2).

### Data Preprocessing

Microarray data preprocessing was performed with the “affy” R package using the “rma” function with default parameters. We created a gene expression table by choosing, for each gene, the probe-set with maximal mean value across all arrays using the “collapseRows” R function from the WGCNA package.

After performing PCA, three of the samples (GSM1287965, GSM1287967, and GSM1287968) were observed to be clear outliers in the latent space formed by the first three principal components. Since the heterogeneity that these tumors represent cannot be effectively modeled with only three samples, combined with the fact that the algorithms we employed might be susceptible to these outliers, we decided to remove these samples from the rest of the analysis.

### Data visualization and clustering

PCA was performed in Matlab using the “pca” function with the default SVD algorithm. For hierarchical clustering, we used the Matlab function “clustergram” with standardized rows (=genes or features), Euclidean distance metric, and average linkage otherwise specified.

### Archetype analysis

The archetypes and best fitting simplex containing the data points (=tumors) were calculated using the “ParTI_lite” matlab function (https://www.weizmann.ac.il/mcb/UriAlon/download/ParTI) with default parameters. Pareto Task Inference (ParTI), is a method for inferring biological tasks from high dimensional data [11]. The function finds the shape of the best fitting polytope (triangle, tetrahedron, etc.) which encompasses the data points. The vertices of this polytope, or “archetypes”, represent biological tasks, and data points specialize at each task according to their distance from the archetypes. The identity of the tasks can be inferred from features enriched near the archetypes.

For Gene Ontology Enrichment analysis, lists of genes found to be over-expressed in each of the archetypes were used as inputs to ToppGene [15]. Venn diagrams were prepared using the matplotlib function “Venn2” in python.

### Topic Modeling

“Topic models” or “grade of membership models” are used in natural language processing to model documents that contain words from different “topics” [16]. Given a set of documents and an assumed number of topics k from which they are composed, it is possible to infer the best fitting parameters of the model by using the Expectation Maximization (EM) algorithm, and thus discover both the k latent topics as well as their proportions in each individual document. Other applications are in population genetics to model individuals with mixed ancestry, and in gene expression datasets to model samples with partial memberships in multiple biologically-distinct clusters [17].

In this study we used the Latent Dirichlet allocation (LDA) model. This model assumes that the given set of documents can be characterized by a Dirichlet distribution with concentration parameters *α*_1_*, α*_2_, …*, α*_*k*_. Each individual document in the set is described as a mixture of k latent topics with proportions described by the random numbers *θ*_1_*, θ*_2_, …*, θ*_*k*_ (whose sum equals to one) that are chosen from the Dirichlet distribution. Each word in the document is independently generated by first selecting one of these topics (according to the probabilities *θ*_1_*, θ*_2_, …*, θ*_*k*_) and then sampling a word from the dictionary (that is, the distribution over words) associated with the chosen topic.

In our case, each document corresponds to a tumor, each topic corresponds to an “idealized” tumor, each word in the vocabulary (“word bag”) corresponds to a gene, and the number of occurrences of a specific word in given document (or topic) corresponds to the expression level of that specific gene in the specific tumor (or “idealized” tumor). Thus, fitting a topic model to a dataset of gene expression profiles from a set of tumors enables us to infer the latent topics (that presumably represent “idealized” tumors) and also the proportions of topics from which every single tumor is composed.

The parameters of the topic model were learned using the “fit_topic_model” function from the R package “fastTopics” [13] with the default number of iterations. The number of latent topics was set between k=2,3,…,10. The outputs of the learning algorithm are: (1) The “Loadings” L matrix which contains the proportions of the k topics *θ*_1_*, θ*_2_, …*, θ*_*k*_ in each tumor; (2) The “Factors” F matrix which contains the distributions over genes *p(gene*|*topic)* for each of the k topics. Log fold changes between topics were computed using the fastTopics function “de_analysis”. To calculate the posterior probabilities *p(gene*|*topic)* we used Bayes’ theorem:

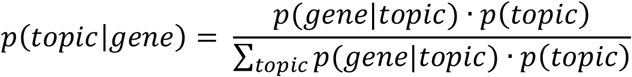

Where we set the prior *p(topic)* = 1/3 for each topic.

Choosing the number of clusters in unsupervised learning problems is often a thorny issue. Methods for choosing k do exist: the maptpx R package [18], for example, computes Bayes Factors for model fits. In general, however, there is no one “true” value, and sometimes different values of k can even complement each other [19]. In our case we chose k=3 topics since the tumors form a triangle in latent space which indicates that they can be modeled as mixtures over three topics. To test the robustness of our analysis, we repeated it with the maptpx R package [18] and obtained similar results.

### Cellular deconvolution

Cellular deconvolution takes a “bulk” gene expression measurement, along with a reference single-cell gene expression matrix and its cell-state space (e.g. a 2D projection such as PCA or tSNE). The algorithm then randomly samples a representative subset of cells from the single-cell matrix and performs a “deconvolution” step, in which the single-cell composition of the bulk sample is inferred using support vector regression (SVR). The process is repeated to ensure that all single cells will be chosen a sufficient number of times, and the inferred abundances are then averaged over all iterations and smoothed over the cell-state space using a pre-selected number of nearest neighbors.

In this study we used the Cell Population Mapping (CPM) R package [14]. For the single-cell reference matrix we used the transcriptomes of 544 individual cells previously collected from a mouse fetal kidney [20]. tSNE was done using MATLAB with default parameters.

## RESULTS

### Pareto Task Inference shows that blastemal type, post-operative chemotherapy Wilms’ tumors fill a triangle shaped continuum in latent space that is bounded by archetypes with stromal, epithelial, and blastemal characteristics

We first downloaded a dataset containing gene expression microarray measurements from 53 high-risk, blastemal type, post-operative chemotherapy Wilms’ tumors that were published by Wegert *et al*. [8]. After preprocessing and data standardization, we performed principal components analysis (PCA) and found that the tumors are arranged in a triangle-shaped continuum in latent space, rather than discrete separated clusters (Figure 1A-F). Following the principle of Pareto Task Inference [11,21], we assume that the vertexes of this triangle-shaped geometric configuration represent “idealized” tumor components or cellular “archetypes” from which all the tumors within the triangle are composed. We therefore used the ParTI MATLAB package developed by Alon and colleagues [11,22] to find these archetypes, which are the vertexes of the optimal fitting triangle encompassing the tumors in our dataset. The archetypes were then presented in the same space as the original data points (Figure 1A-F).

In order to characterize and identify the three archetypes, we first chose a set of genes that mark the three main lineages in the developing kidney (the un-induced mesenchyme, the Cap mesenchyme, and early renal epithelial structures) and examined their pattern of expression in latent space (Figures 1A-C). We found that FN1, a gene that is known to be highly expressed in the un-induced mesenchyme, is highly expressed near the first archetype. The membrane transporter gene SLC12A1, that is known to mark epithelial tubules in the developing kidney, is over expressed near the second archetype. The gene SIX2, which marks the Cap mesenchyme, is highly expressed near the third archetype. We therefore labeled the three archetypes as “stromal”, “epithelial”, and “blastemal” respectively. Moreover, we found that TOP2A and MKI67, genes that are known to be over-expressed in proliferating cells, are highly expressed near the blastemal archetype (Figures 1D-E), which is consistent with the proliferative and “aggressive” nature of blastemal tumors after chemotherapy [8].

To confirm our characterization of the three archetypes, we selected a set of 102 genes (Table S3) that are known from the literature to mark specific cell populations in the developing kidney. Then, we performed hierarchical clustering of the tumors in our dataset and their archetypes with respect to these genes (Figure 1G and Figures S1-S12). Indeed, it can be seen that markers for the un-induced mesenchyme (COL1A1, COL3A1, COL5A2, FN1, and SERPINE1) are over-expressed in the stromal archetype, markers of the Cap mesenchyme (SIX2, CITED1, EYA1, and SALL2) are over-expressed in the blastemal archetype, and markers for the renal epithelial tubules (SLC12A1, AQP2, LRP2, and UMOD) are over-expressed in the epithelial archetype. We noticed that the stromal archetype also over-expresses genes characteristic to immune cells (C1QA, C1QB, CCL2, CD14, CD93, and CXCL2), which is consistent the fact that the un-induced mesenchyme contains a relatively large number of infiltrating immune cells (e.g. macrophages). We also note that the stromal archetype over-expresses genes characteristic to muscle cells (DES, MYL1, MYH3, and MYOG), which is consistent with the fact that some stromal tumors have been known to contain muscle-like cells.

We further characterized the archetypes using GO enrichment analysis (Figure 2). For each of the three archetypes, we found a list of genes that are over-expressed (log2FC>2) with respect to both of the other two archetypes (Figure 2A-C, Tables S4-S6). We then inserted the three lists if genes into Toppgene [15]. We found that the stromal archetype is enriched for genes typical to the un-induced mesenchyme, for example, components of the extracellular matrix (Figure 2D). The blastemal archetype is enriched for genes involved in maintenance of nephron progenitors and the mesenchymal to epithelial transition (MET), which are characteristic of the Cap mesenchyme. Finally, genes enriched in the epithelial archetype are involved in cell-cell junctions, transport, a renal epithelial differentiation, which are typical to the early epithelial structures in the developing kidney.

We also checked the relation between the reported tumor histology and clinical parameters to its location in latent space (Figure 1F, Figs S13-S23). We observed that tumors with anaplastic histology (diffuse or focal), which is considered least favorable [8], as well as tumors with mutations in the genes SIX1, SIX2, or DROSHA, tend to cluster in the vicinity of the blastemal archetype. This agrees with the higher incidence of SIX1/2 mutations in tumors with chemotherapy-resistant blastema that was observed by Wegert *et al*. [8]. Likewise, we observed that the single blastema-only xenograft in the dataset is also located closest to the blastemal archetype more than any other tumor. This is consistent with previous observations that patient-derived xenografts significantly increase the percentage of their blastemal component from their first passage [23].

### Topic modeling shows that each tumor can be represented as a unique mixture of three hidden topics with blastemal, stromal, and epithelial characteristics

The fact that the tumors in our dataset create a triangle-shaped continuum in latent space suggests that each tumor can be represented as a unique mixture of three “idealized” tumor components. Therefore, in order to provide a more quantitative interpretation, we fitted a topic model with k=3 hidden “topics” to our dataset [13] (see Methods). This allowed us to infer both the three latent topics (that presumably represent the “idealized” tumor components) and also the proportions of topics from which every single tumor is composed. We observed that, indeed, tumors located near each of the three vertexes of the triangle-shaped continuum are predominantly composed one out of the three topics (Figure 3A), whereas tumors located in-between the vertexes of the triangle or near its center are composed of multiple topics (Figures S24-S25 and Supplementary data).

We next set to identify the three topics (Figure S26). We observed that topic no. 1 over-expresses markers for the renal epithelial tubules (e.g. CDH1, SLC12A1, LRP2, and UMOD) and we therefore labeled it the “epithelial” topic. Likewise, topic no. 2 over-expressed markers for the un-induced mesenchyme (e.g. COL1A1, COL1A2, TWIST1, ZEB2, and SERPINE1) and we therefore labeled it the “stromal” topic, and topic no. 3 over-expressed markers for the Cap mesenchyme (e.g. SIX2, CITED1, EYA1, and SALL2) and we therefore labeled it the “blastemal” topic.

We showed this also by calculating the posterior probabilities *p(gene*|*topic)* over all the genes for each of the three topics (Figure 3B). The posterior probability of a specific topic given the expression of a specific gene represents the association between over-expression of that specific gene and the probability of that specific topic being represented in the tumors of our dataset. Indeed, we observed that that over-expression of markers for the renal epithelial tubules (e.g. CDH1, SLC12A1, LRP2, and UMOD) is associated with high probability for the epithelial topic (topic no. 1), over-expression of markers for the un-induced mesenchyme (COL1A1, COL1A2, COL5A2, FN1, and SERPINE1) is associated with high probability for the stromal topic (topic no. 2), and over-expression of markers for the Cap mesenchyme (SIX2, CITED1, EYA1, and SALL2) is associated with high probability for the blastemal topic (topic no. 3). To further confirm this, we selected a list of genes characterizing each topic, that is, genes for which the posterior probability *p(gene*|*topic)* > 0.5. Using GO enrichment analysis as before, we found that indeed, the three hidden topics over-express genes related to epithelial (resembling early renal tubular epithelium), stromal (un-induced mesenchyme-like), or blastemal (Cap mesenchyme-like) cell types (Tables S7-S9).

We also checked the relation between the topic composition of each tumor to its histology and clinical parameters (Figure 3C and Figure S27). We found that tumors with anaplastic histology (diffuse or focal), which is considered least favorable, as well as the single blastemal xenograft in the dataset, and also tumors with mutations in the genes SIX1, SIX2, or DROSHA, all contain a large fraction of the blastemal topic.

### Cellular deconvolution indicates that each tumor is composed of a unique mixture of cell populations resembling the un-induced mesenchyme, the cap mesenchyme, and the early epithelial structures of the fetal kidney

We next used cellular deconvolution [14] to infer a more detailed cellular composition of each tumor. Since cells in Wilms’ tumors closely resemble those of the fetal kidney, we used a previously published single-cell gene expression dataset from a fetal mouse developing kidney [20] as reference. The cellular deconvolution algorithm provided a prediction of the proportions of ten cell types from the developing kidney within each of the tumors and archetypes (Figure 4 and Figure S28).

We observed that the stromal archetype is composed primarily of cells resembling the un-induced mesenchyme (UM) and macrophages, the epithelial archetype is composed mainly of epithelial cells resembling those of the fetal proximal tubule (PROX_2), the Loop of Henle (LOH), and the distal tubule (DIST_CD), and the blastemal archetype is composed primarily of cells resembling the Cap Mesenchyme (CM), along with some cells resembling very early epithelial tubular structures (PROX_1). Tumors that are near the archetypes in latent space are likewise composed primarily from these cell types, while tumors in between the archetypes or in the middle of the triangle shaped continuum are composed of more heterogeneous mixtures of the different cell populations (Figure S28 and Supplementary data).

We also correlated the reported histology with the cell type repertoire inferred by cellular deconvolution (Figure 4C and Figure S29). We found that most of the tumors with reported blastemal histology contain a significant proportion of cells resembling those of the Cap mesenchyme (CM_ALL), as expected. Likewise, tumors with anaplastic histology (diffuse or focal), as well as the single blastemal xenograft in our dataset, and also tumors reported to contain mutations in the genes SIX1, SIX2, or DROSHA, contained a significant fraction of cells resembling the cycling Cap mesenchyme cells (CM_DIV) in the fetal kidney. This is in agreement with the findings of Wegert *et al*. [8] that blastemal-type Wilms tumors with mutations in SIX1 or SIX2 have a gene expression signature of proliferation and kidney progenitors. On the other hand, tumors reported to have triphasic/mixed or regressive histology were found to contain high proportions of cells resembling the un-induced mesenchyme (UM).

## DISCUSSION

The dataset used in this study was collected by Wegert *et al*. [8] from high-risk, blastemal type, post-operative chemotherapy Wilms’ tumors that were first treated with chemotherapy, according to the SIOP protocol. We recently reported similar results when performing Pareto task inference in Favorable Histology Wilms’ Tumors (FHWT’s) [10] in microarray datasets collected by the Children’s Oncology Group [24–27]. In that dataset, the microarray measurements were performed on samples of tumors that were surgically resected before chemotherapy, as per the COG protocol. Thus, the triangle-shaped continuum formed by Wilms’ tumors in latent space seems to be a conserved in both datasets (either treated by the COG or by the SIOP protocols), which indicates that it is an intrinsic property of Wilms’ tumors heterogeneity. We note that integration of both datasets is difficult since the studies were performed with microarrays of different types and there is a large technical bias between them. We believe that this problem will be mitigated in the future as more and more tumors are analyzed using RNA sequencing and large scale datasets become publicly available.

## ACKNOWLEDGMENTS

We wish to thank Manfred Gessler, Jenny Wegert, Dudi Feldman, Noam Korngut, Amit Frishberg, Naomi Pode-Shakked, and all members of our lab for helpful comments and suggestions.

## AUTHOR CONTRIBUTIONS

Study initiation and conception – Y.T. and T.K.; Pareto task inference – Y.T. and T.K.; Topic modeling – Y.T., J.G., and T.K.; Cellular deconvolution – Y.T. and T.K.; Other intellectual contribution – A.U., B.D., P.H.; Manuscript writing – Y.T and T.K.

## APPENDICES

**Supplementary information:** Supplementary text and figures.

**Table S1:** Normalized gene expression values of the Wilms’ tumors from Wegert *et al*. and the three archetypes.

**Table S2:** Integrated histological and clinical data from Wegert *et al*., including the mapping from the sample IDs of the mRNA microarrays (WT055, WT0056….) to the sample IDs in the GEO database (dkfz1079, dkfz1080….)

**Table S3:** A list of 102 selected genes that are known from the literature to be associated with kidney development and tumorigenesis.

**Table S4:** Gene Ontology (GO) enrichment analysis for the *stromal* archetype. Genes for which log2FC>2 in the *stromal* archetype with respect to the other two archetypes were selected and inserted to Toppgene.

**Table S5:** Gene Ontology (GO) enrichment analysis for the *epithelial* archetype. Genes for which log2FC>2 in the *epithelial* archetype with respect to the other two archetypes were selected and inserted to Toppgene.

**Table S6:** Gene Ontology (GO) enrichment analysis for the *blastemal* archetype. Genes for which log2FC>2 in the *blastemal* archetype with respect to the other two archetypes were selected and inserted to Toppgene.

**Table S7:** Gene Ontology (GO) enrichment analysis for the epithelial topic (k1). Genes for which the posterior probabilities *p*(*topic* = *k*1|*gene*) > 1/2 were selected and inserted to Toppgene.

**Table S8:** Gene Ontology (GO) enrichment analysis for the stromal topic (k2). Genes for which the posterior probabilities *p*(*topic* = *k*2|*gene*) > 1/2 were selected and inserted to Toppgene.

**Table S9:** Gene Ontology (GO) enrichment analysis for the blastemal topic (k3). Genes for which the posterior probabilities *p*(*topic* = *k*3|*gene*) > 1/2 were selected and inserted to Toppgene.

**Program:** A compressed directory containing programs and datasets for data visualization, and well as sets of figures characterizing each tumor.

