## Supplementary text and figures for "Characterization of the continuous transcriptional heterogeneity in Wilms’ tumors using unsupervised machine learning"

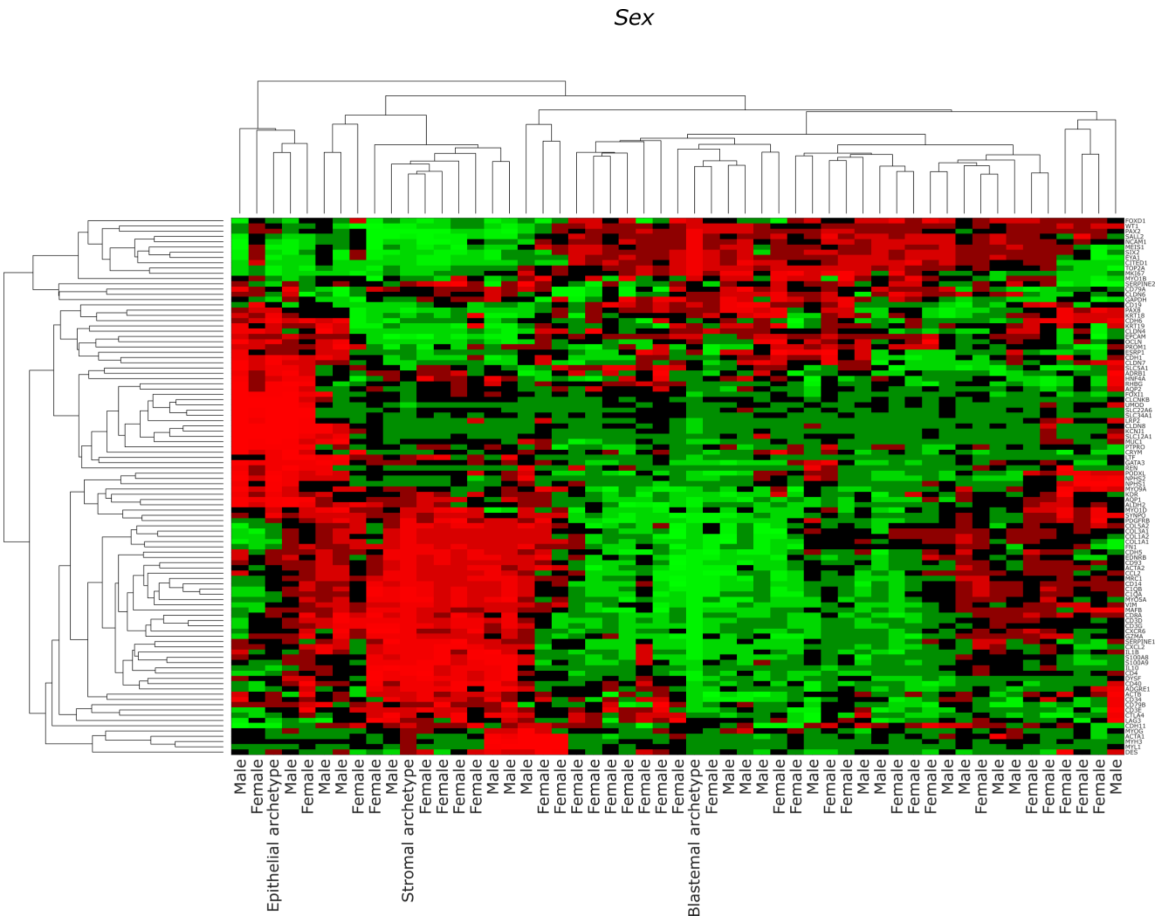


**Figure S1:** A heatmap of selected genes vs. tumors and archetypes showing reported **patient sex** (male/female).

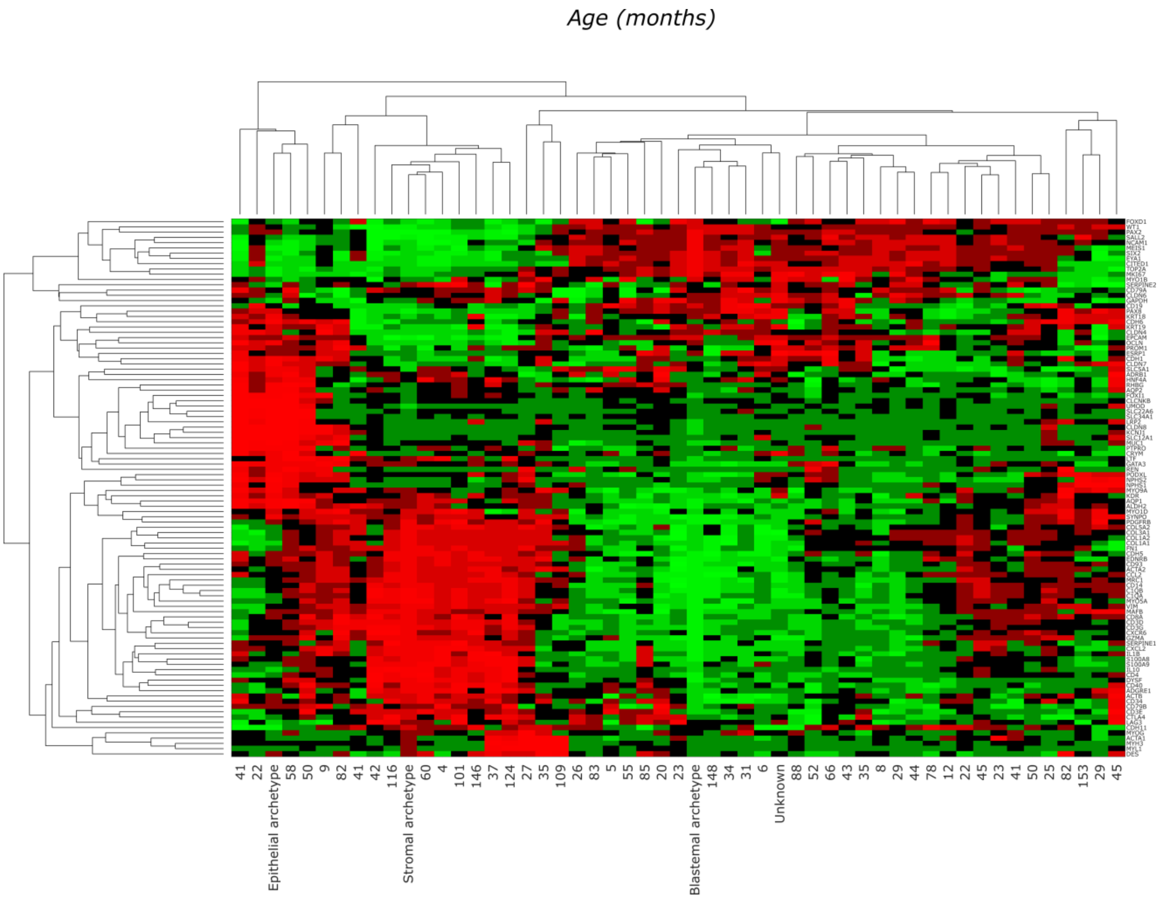


**Figure S2:** A heatmap of selected genes vs. tumors and archetypes showing **patient age** (in months) at diagnosis.

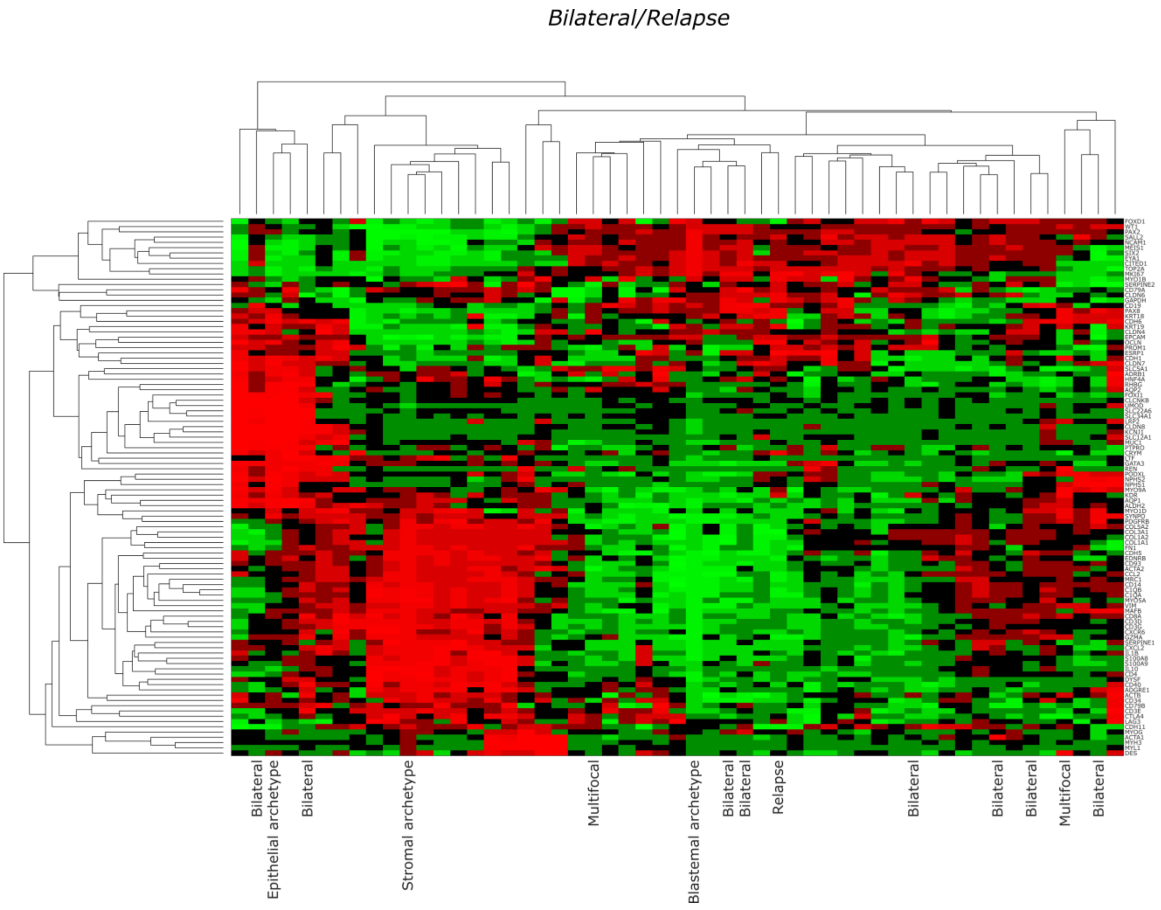


**Figure S3:** A heatmap of selected genes vs. tumors and archetypes showing clinical data (**bilateral and/or relapse**).

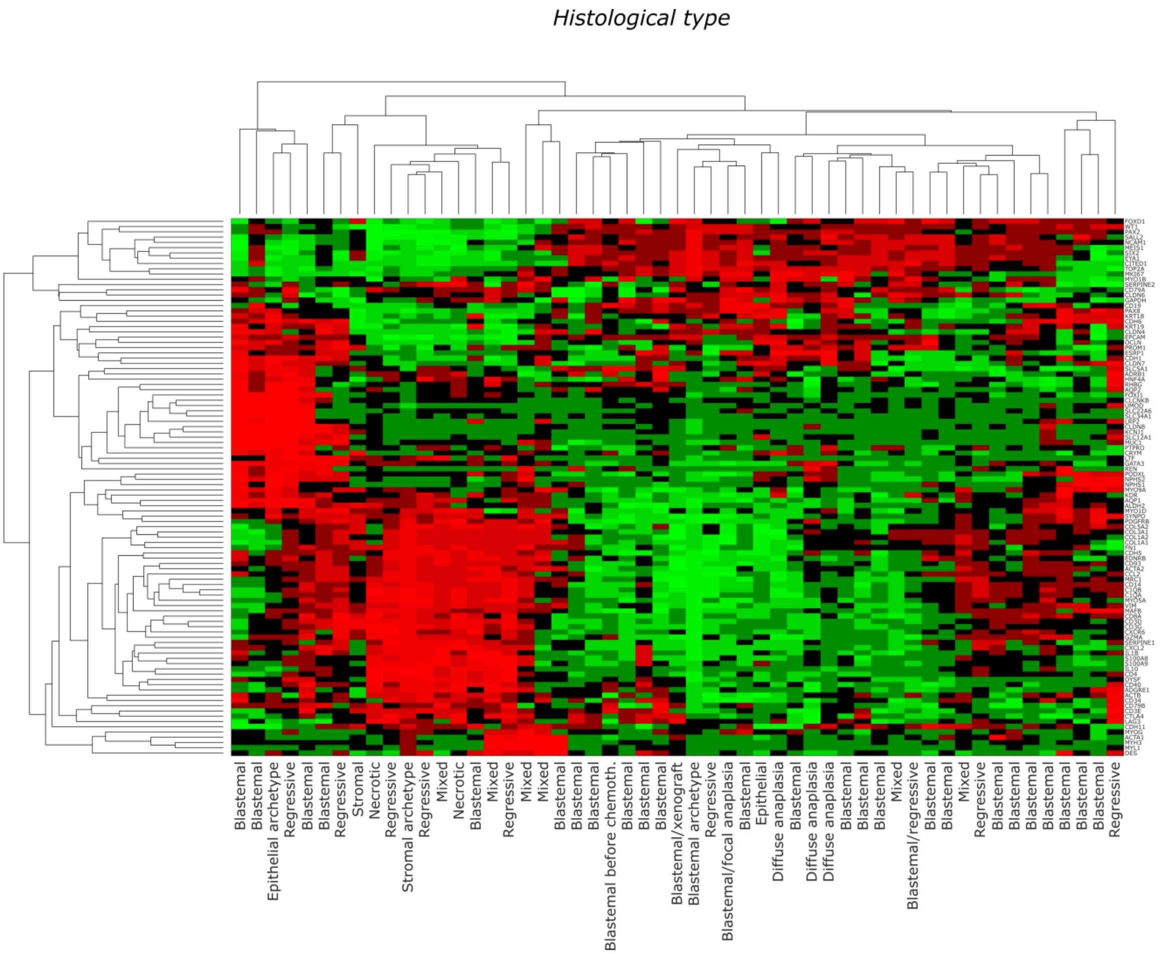


**Figure S4:** A heatmap of selected genes vs. tumors and archetypes showing the histological type.

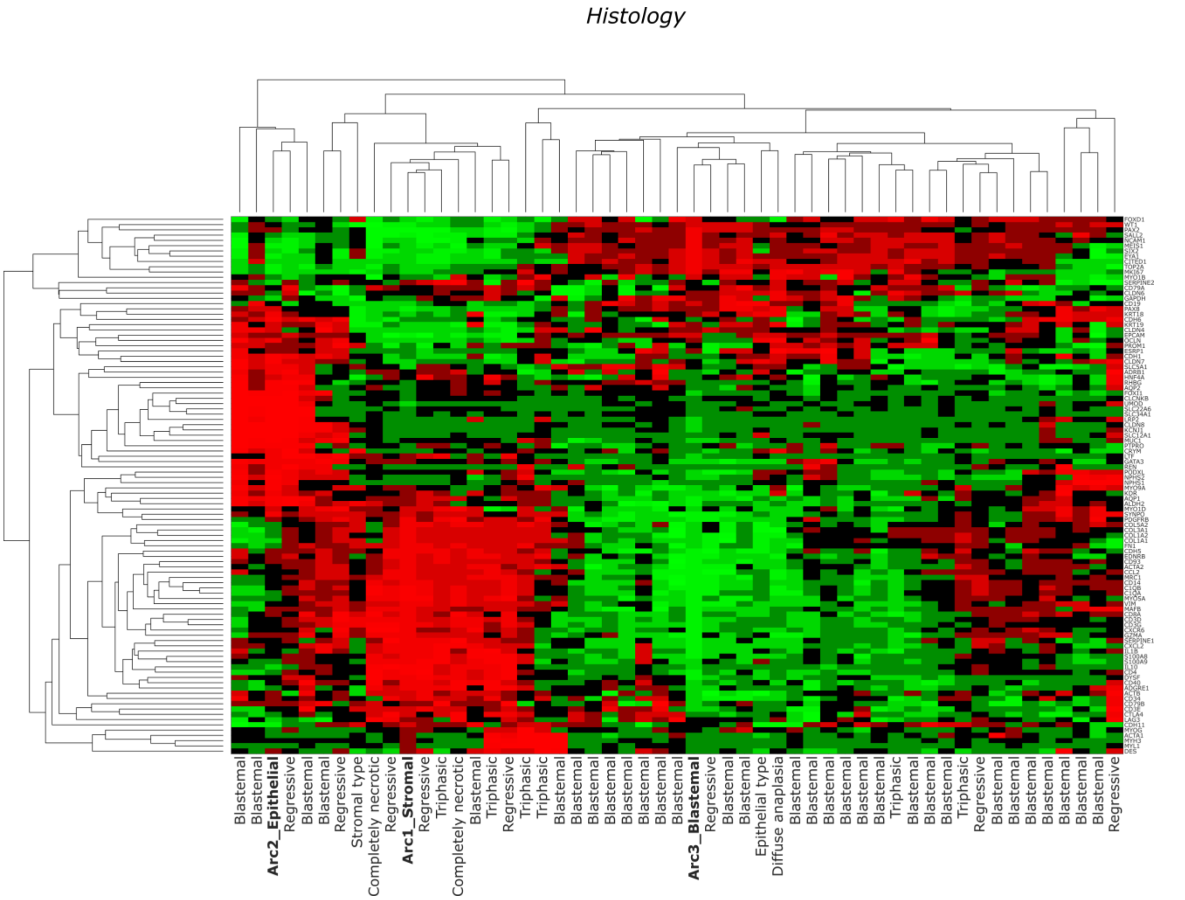


**Figure S5:** A heatmap of selected genes vs. tumors and archetypes showing tumor histology.

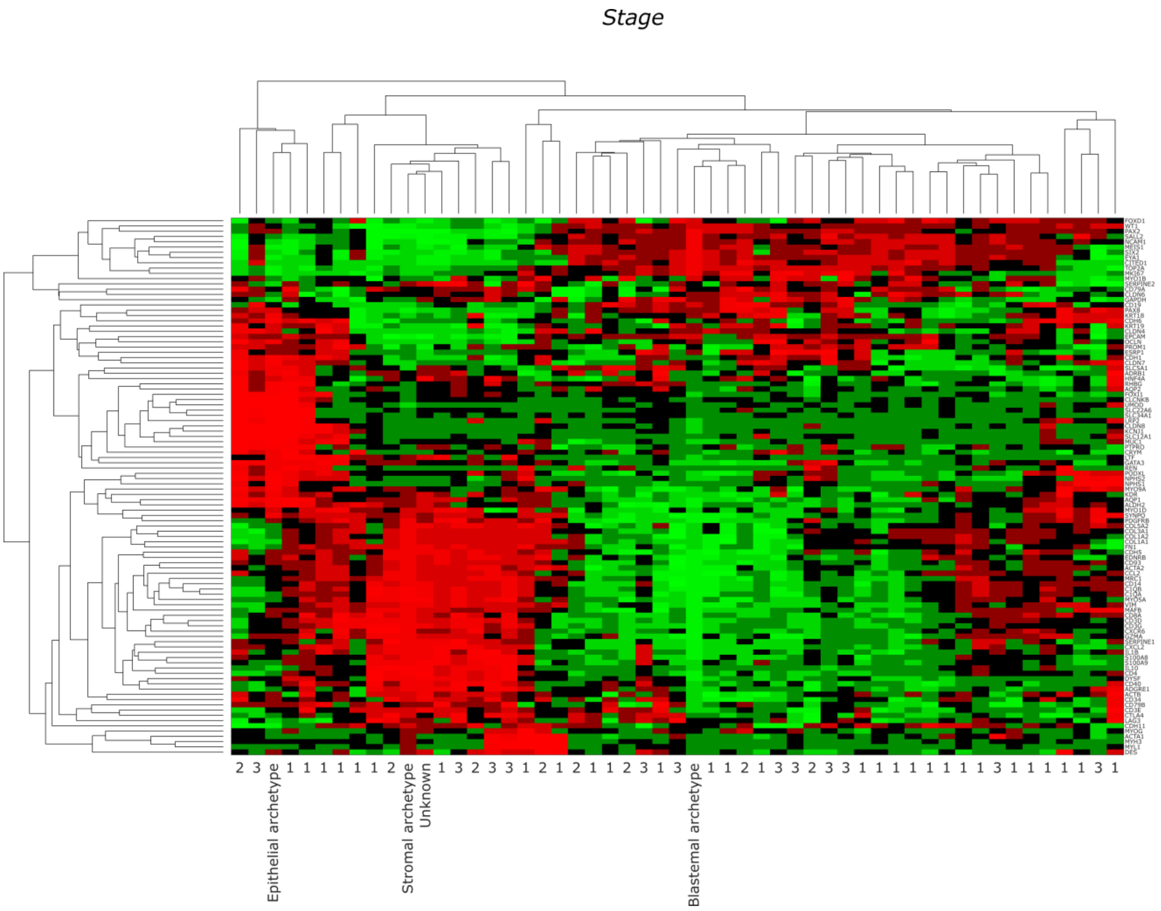


**Figure S6:** A heatmap of selected genes vs. tumors and archetypes showing reported **tumor stage** at diagnosis.

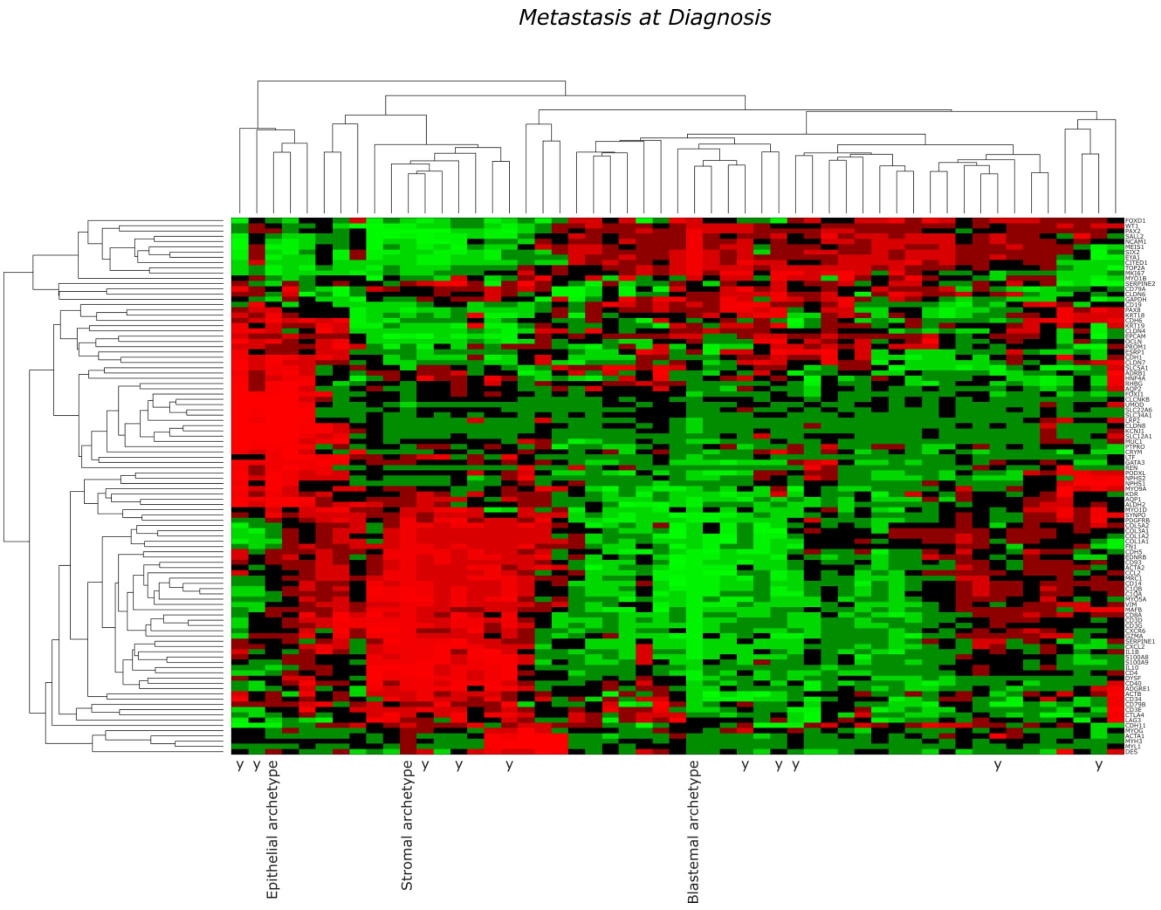


**Figure S7:** A heatmap of selected genes vs. tumors and archetypes showing reported **metastasis at diagnosis**.

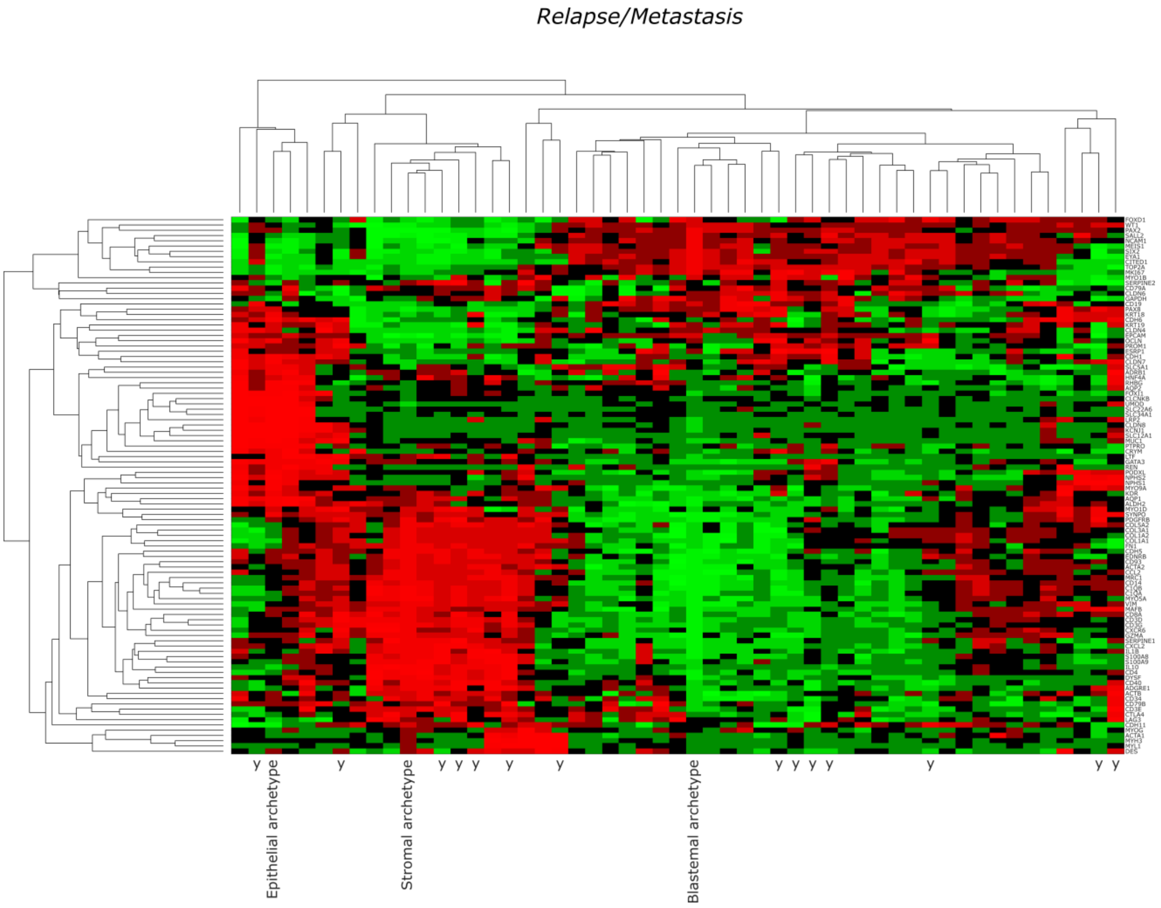


Figure S8: A heatmap of selected genes vs. tumors and archetypes showing the reported status of the tumor (**relapse/metastasis**).

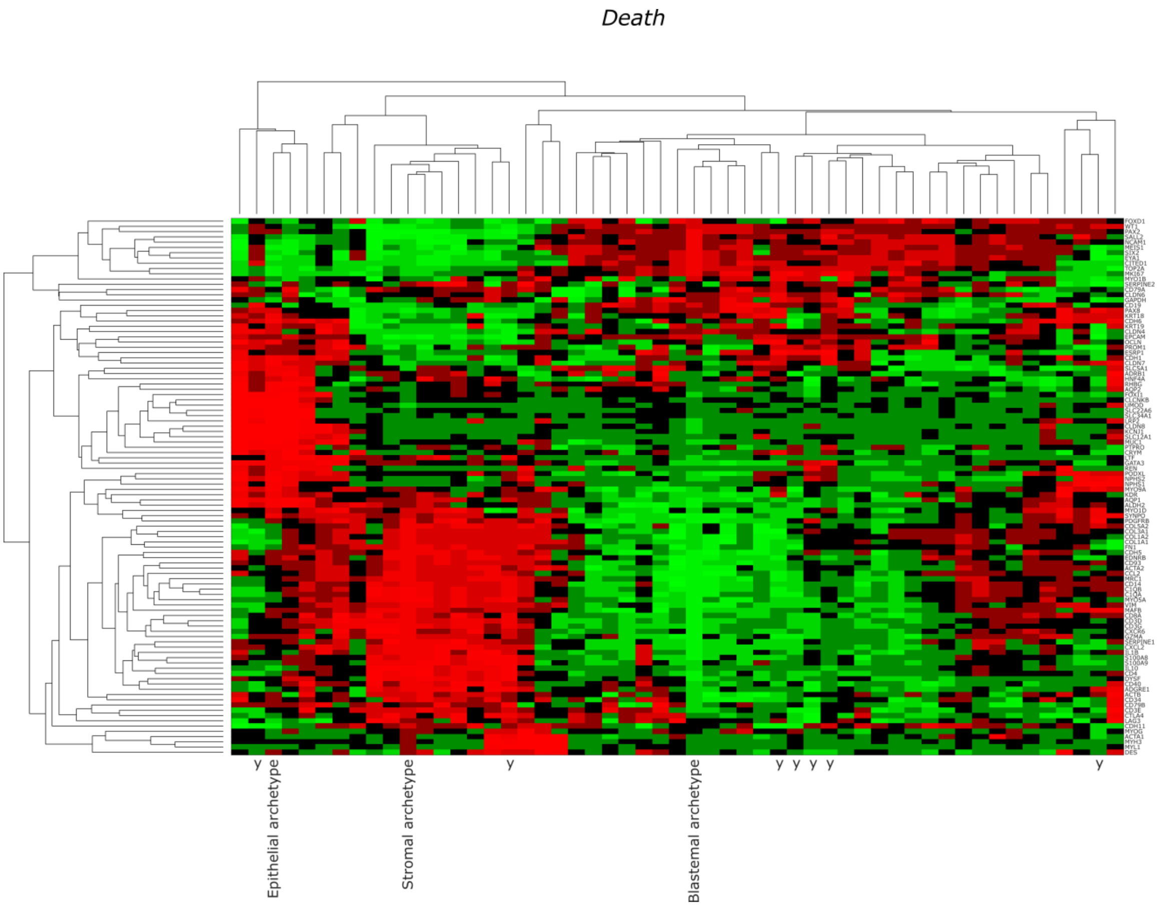


**Figure S9:** A heatmap of selected genes vs. tumors and archetypes showing the reported status of the patient (**death**).

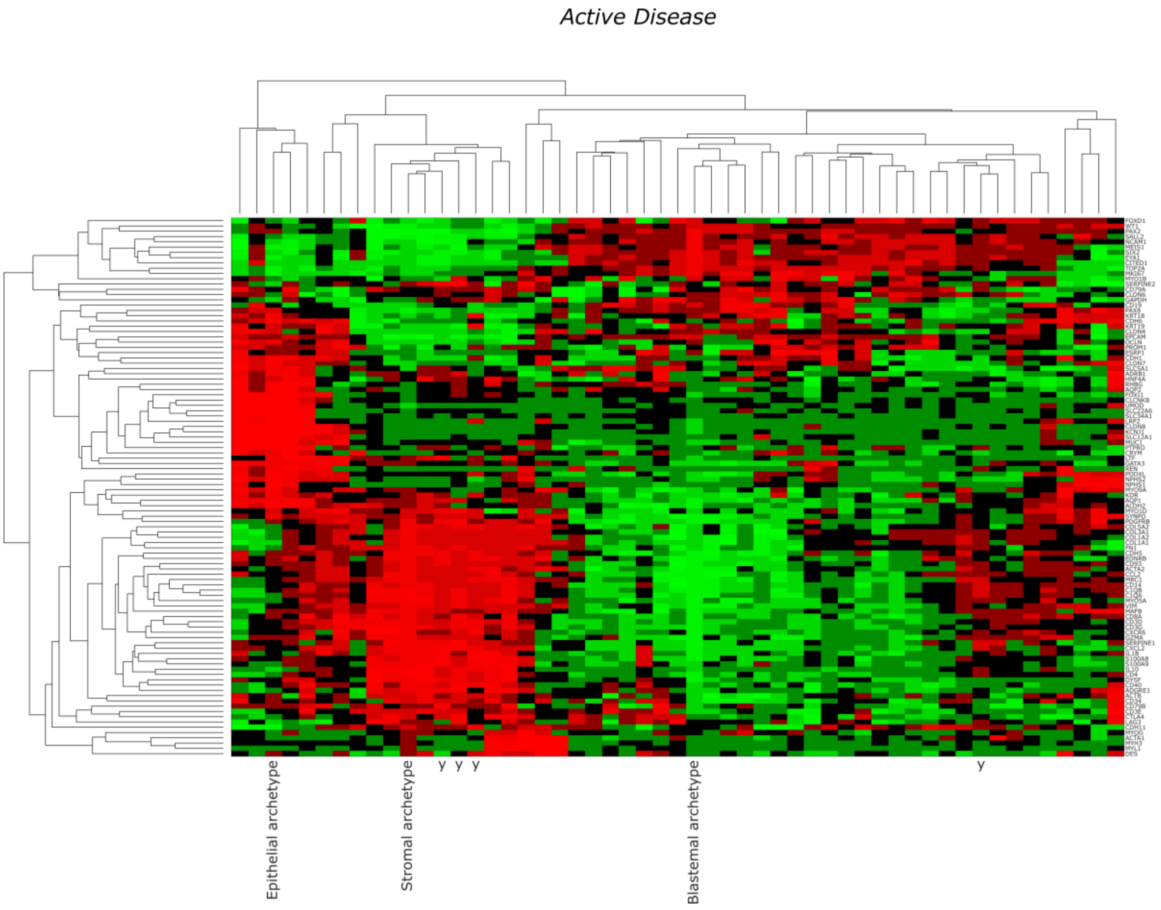


**Figure S10:** A heatmap of selected genes vs. tumors and archetypes showing the reported activity of the tumor (**active disease**). It can be seen that most tumors from patients with active disease in this dataset tend to cluster near the stromal archetype.

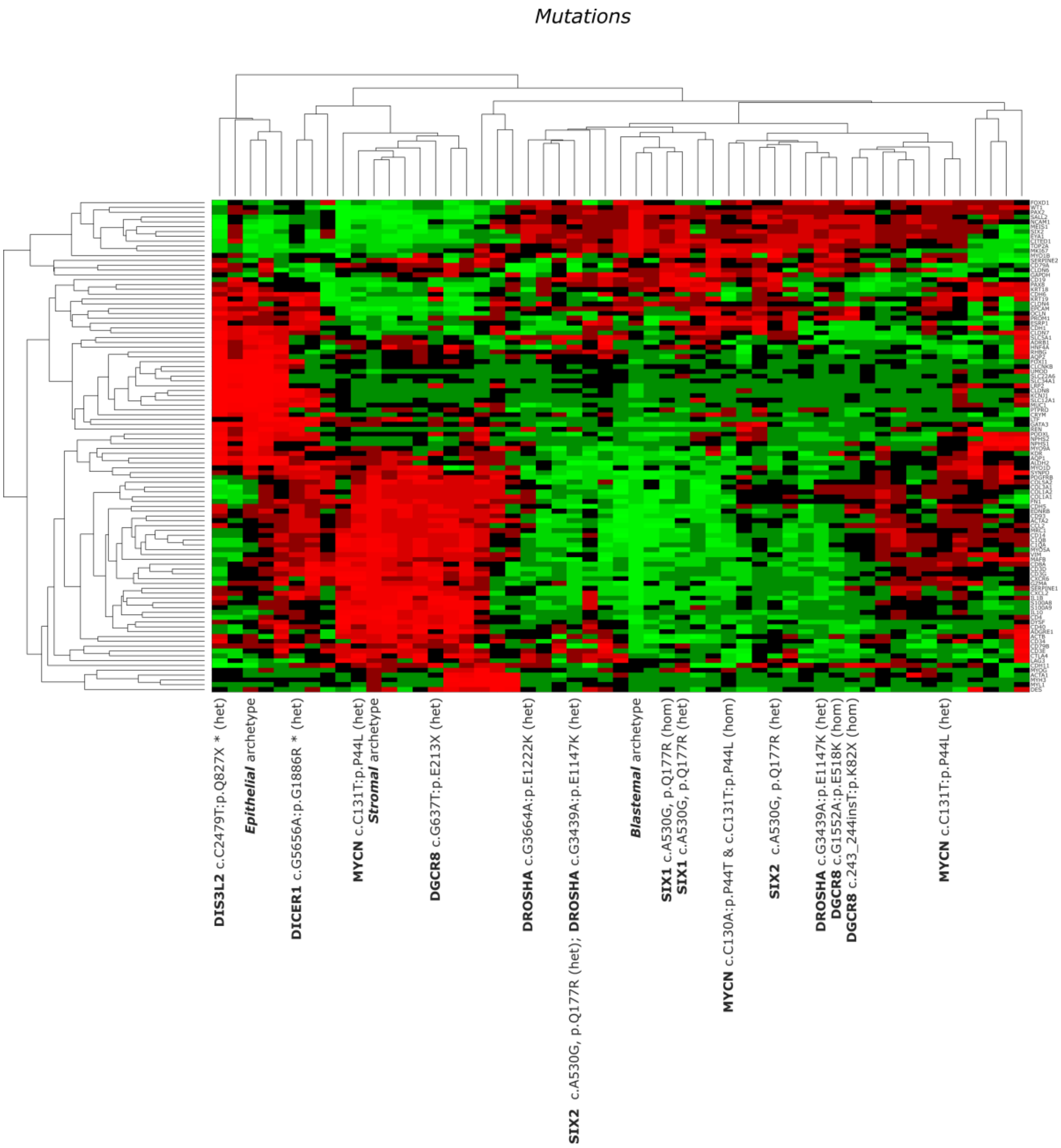


Figure S11: A heatmap of selected genes vs. tumors and archetypes showing reported mutations. It can be seen that mutations in the genes SIX1, SIX2, or DROSHA tend to cluster with the more blastemal tumors.

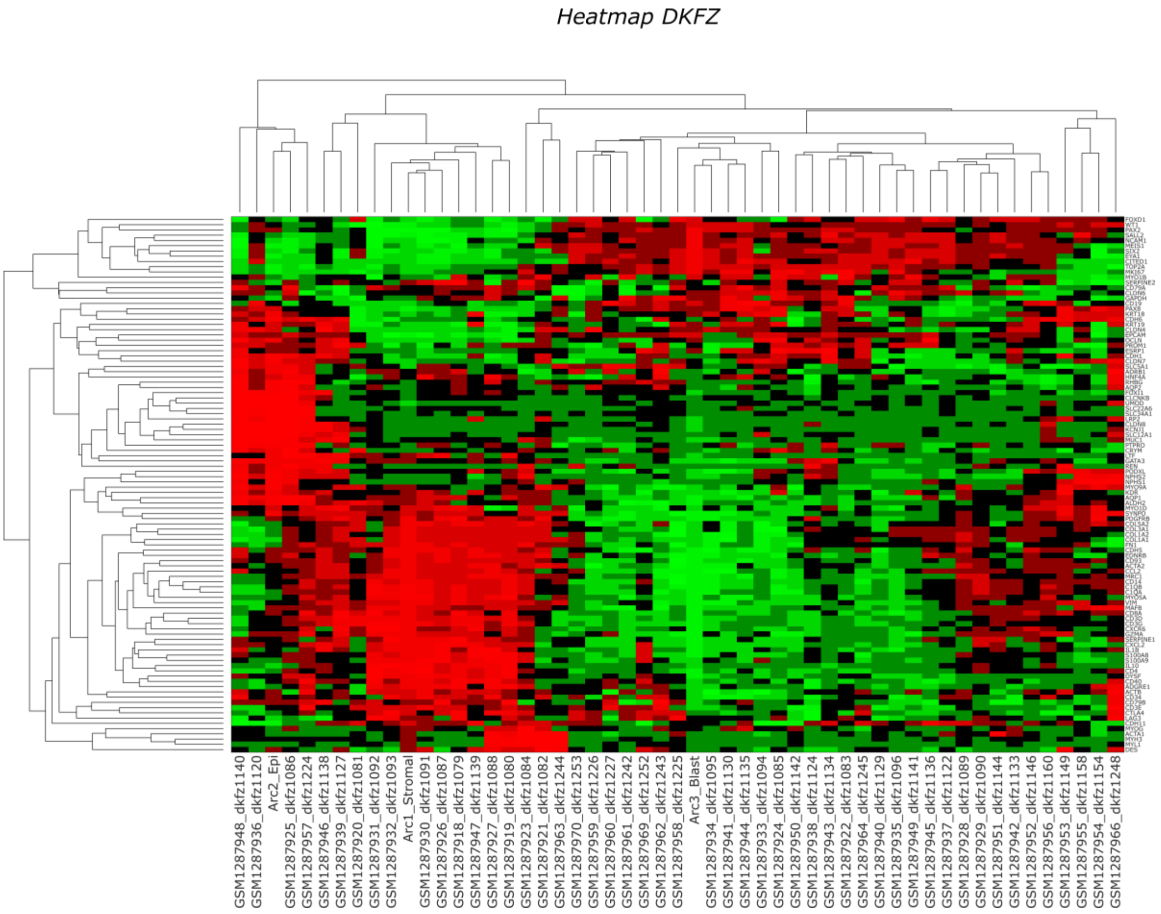


**Figure S12:** A heatmap of selected genes vs. tumors and archetypes showing the **sample ID** (GEO accession number and DKFZ identifier).

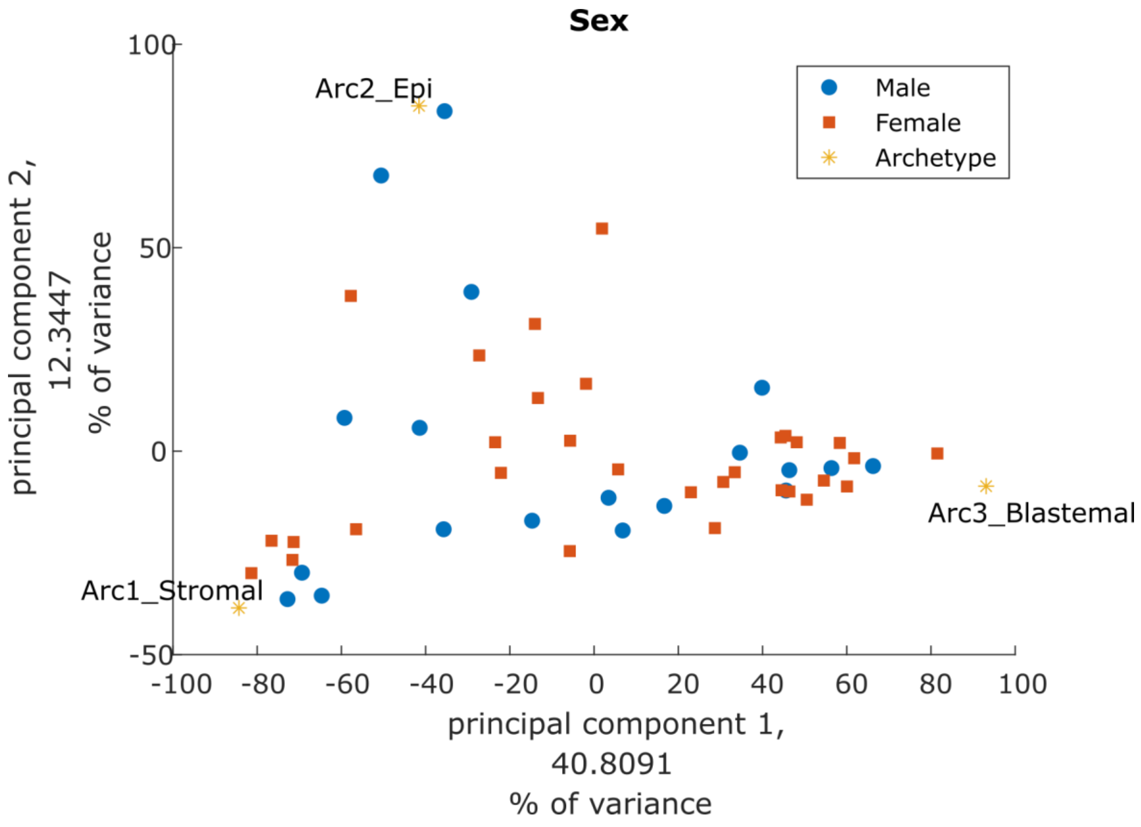


**Figure S13:** PCA of tumors and archetypes showing the reported **patient sex** (male/female).

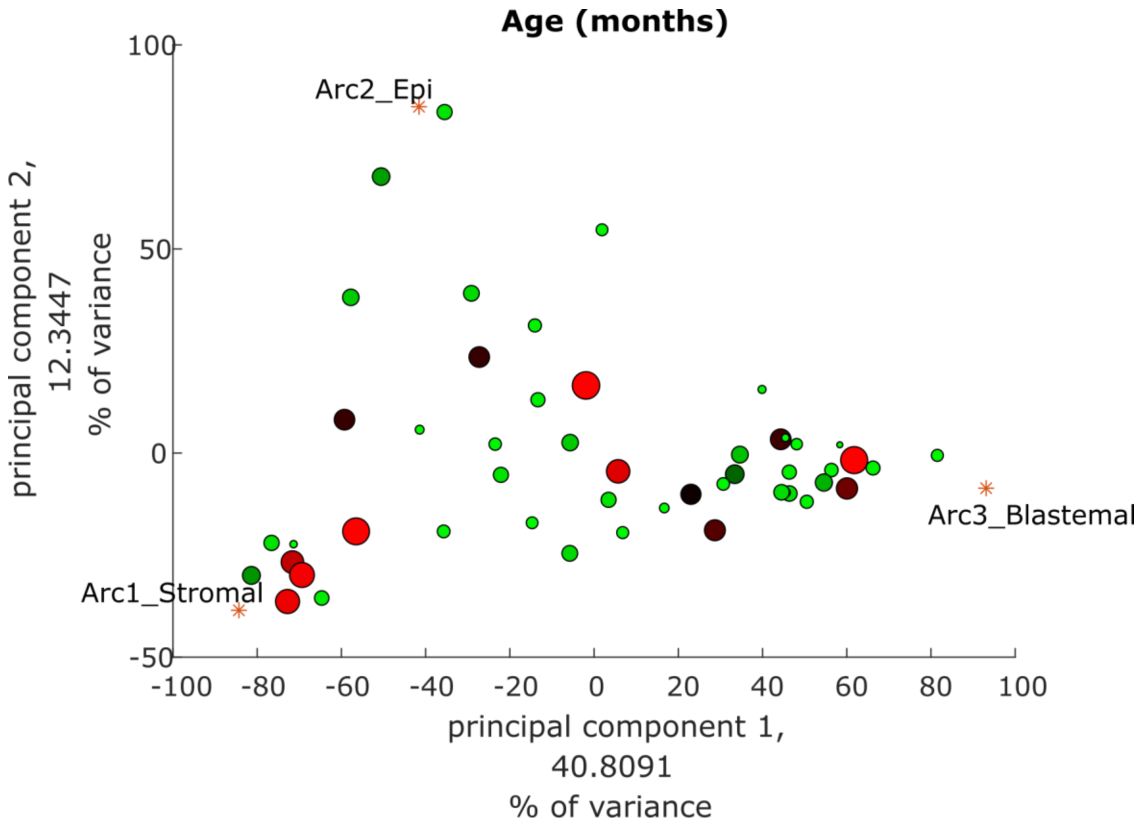


**Figure S14:** PCA of tumors and archetypes showing **patient age** (in months) at diagnosis (old – red/large symbol, young – green/small symbol).

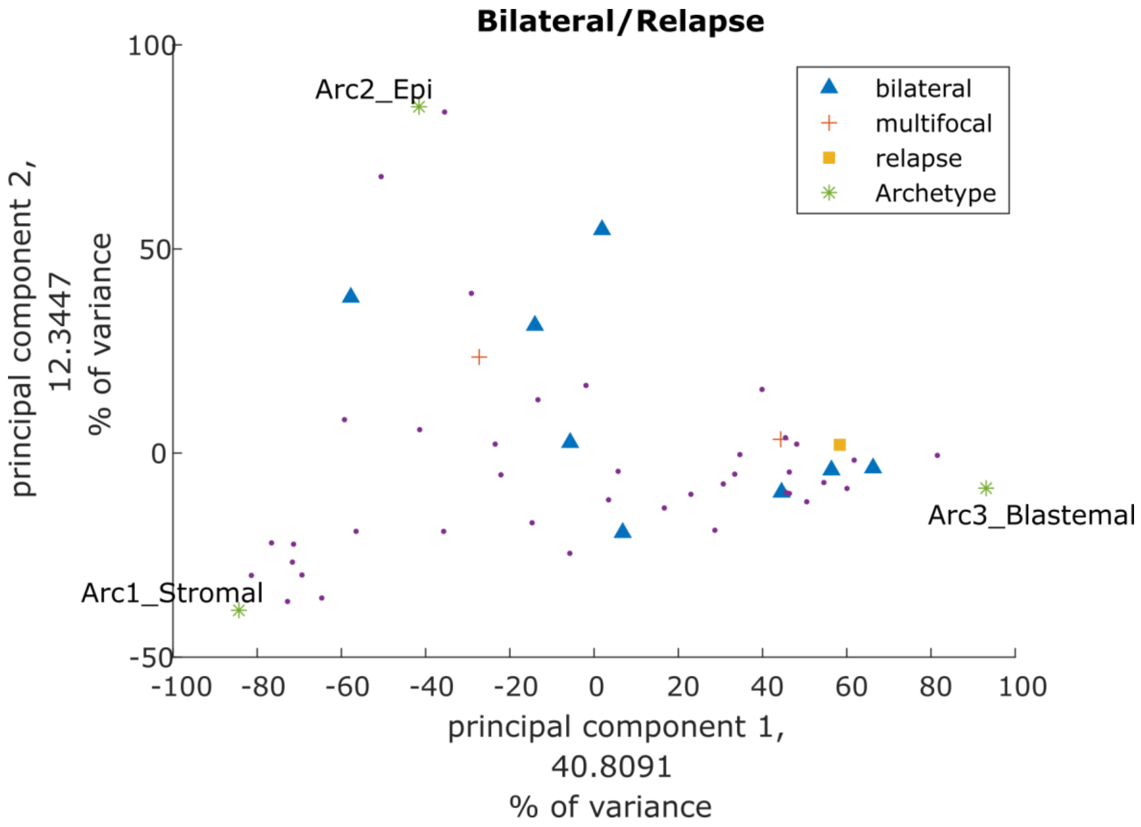


**Figure S15:** PCA of tumors and archetypes showing the reported clinical data (**bilateral and/or relapse**). It can be seen that bilateral tumors tend to cluster away from the stromal archetype.

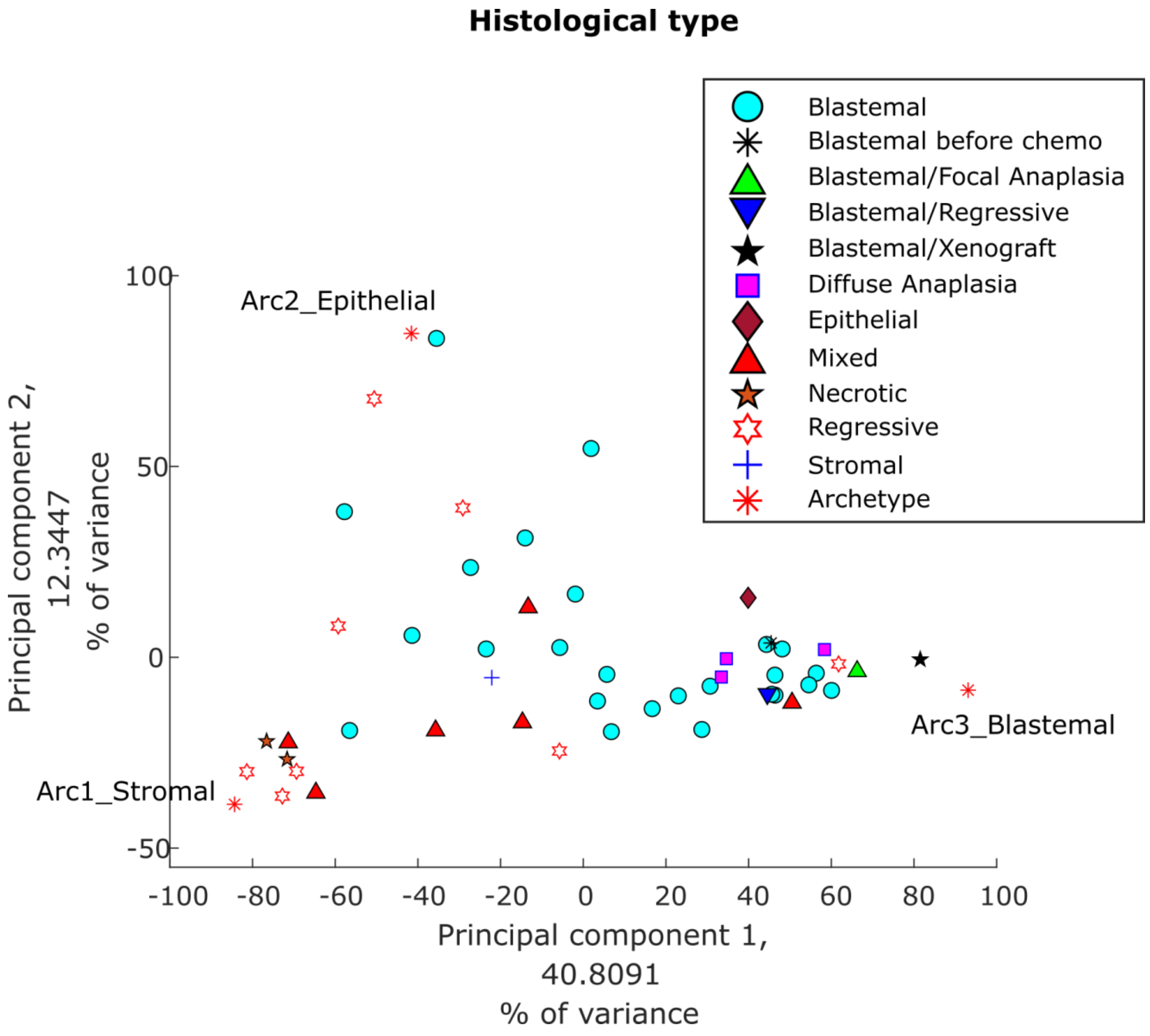


**Figure S16:** PCA of tumors and archetypes showing the reported **histological type**. It can be seen that the tumors with anaplastic histology (diffuse or focal), which is considered least favorable, tend to cluster in the vicinity of the blastemal archetype. Likewise, notice that the single blastema-only xenograft in the dataset is located closest to the blastemal archetype, more than any other tumor. This is consistent with previous observations that patient-derived xenografts significantly increase the percentage of their blastemal component from their first passage [1].

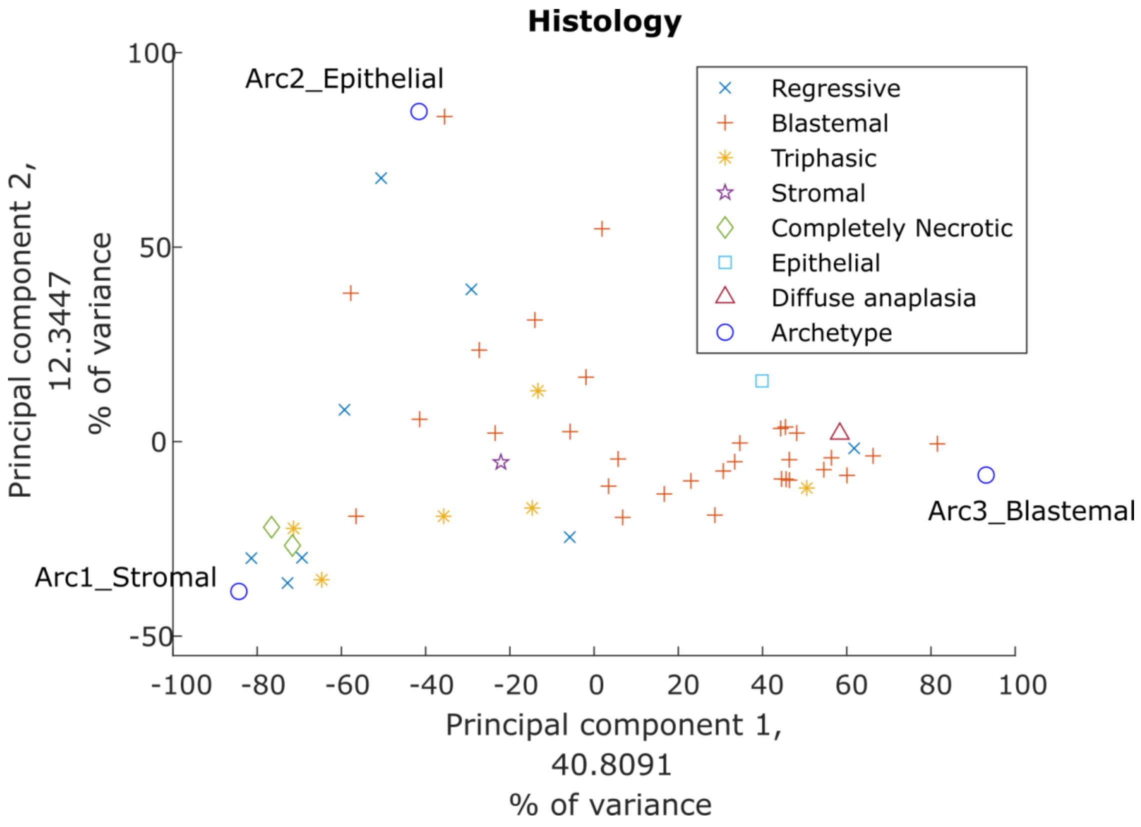


Figure S17: PCA of tumors and archetypes showing the reported tumor **histology**.

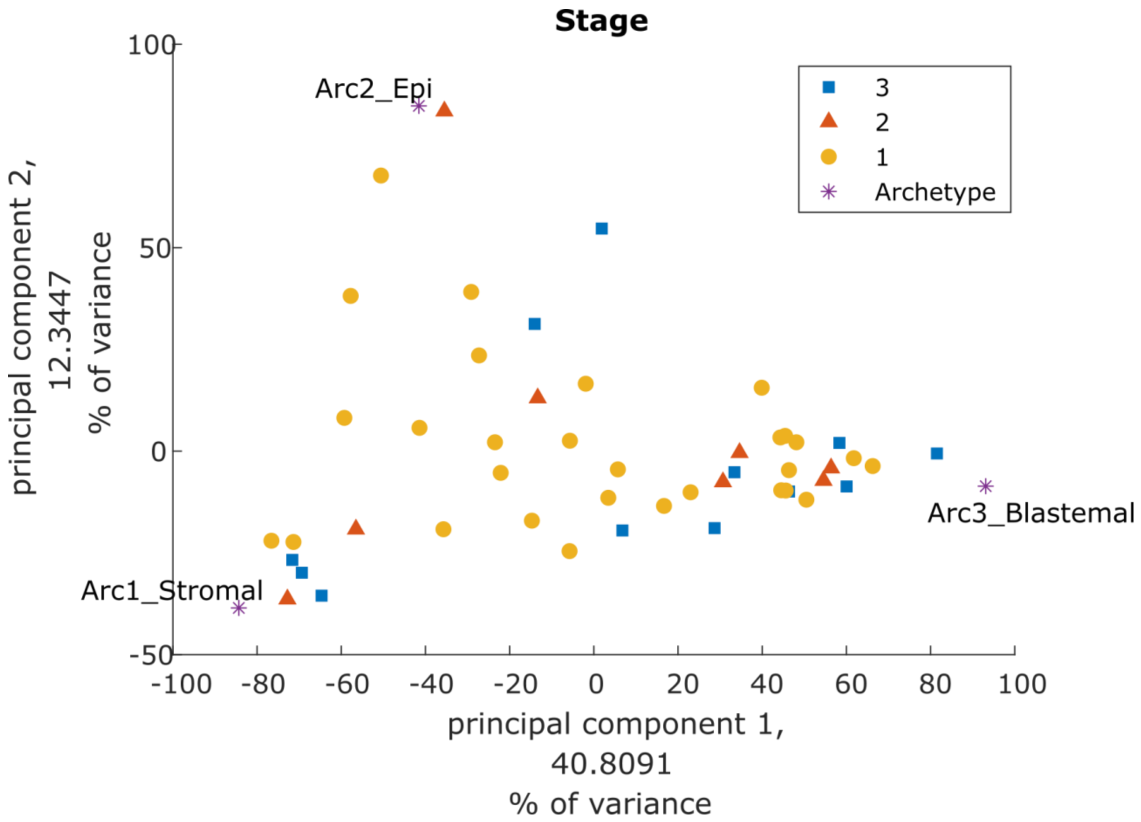


Figure S18: PCA of tumors and archetypes showing the reported tumor **stage** at diagnosis.

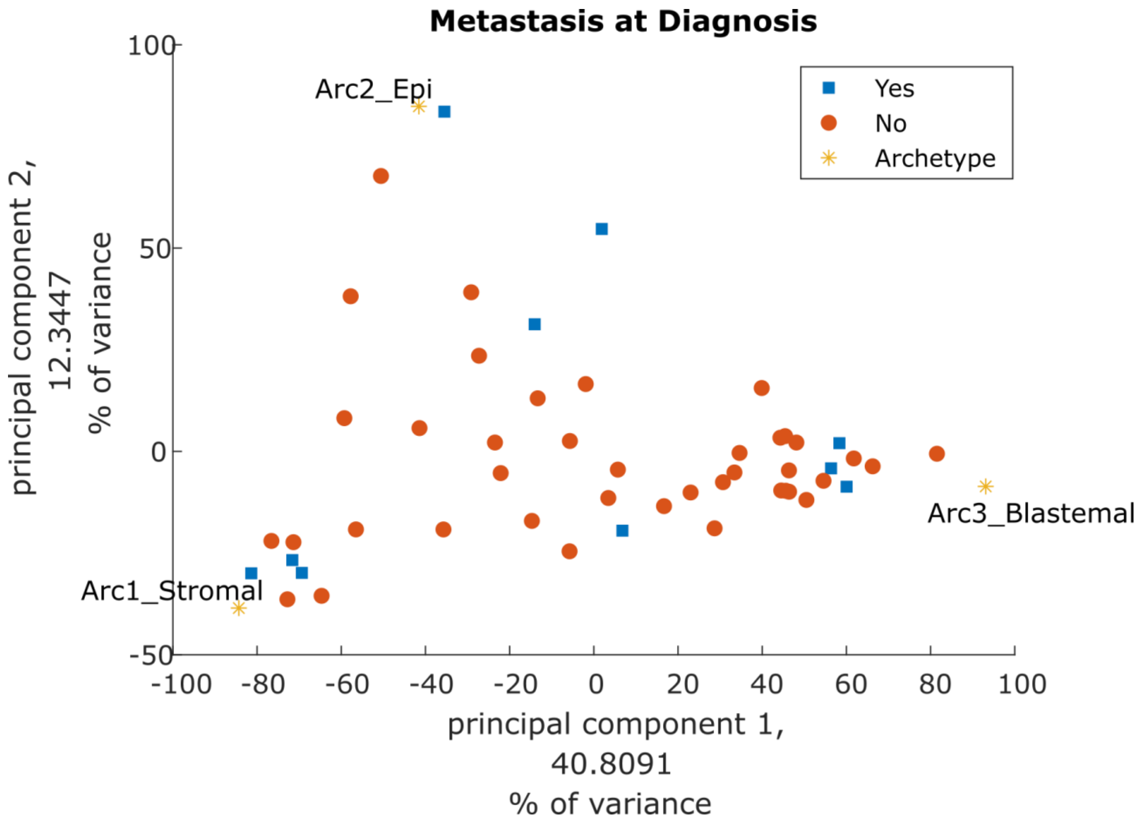


**Figure S19:** PCA of tumors and archetypes showing the reported tumor **metastasis at diagnosis**.

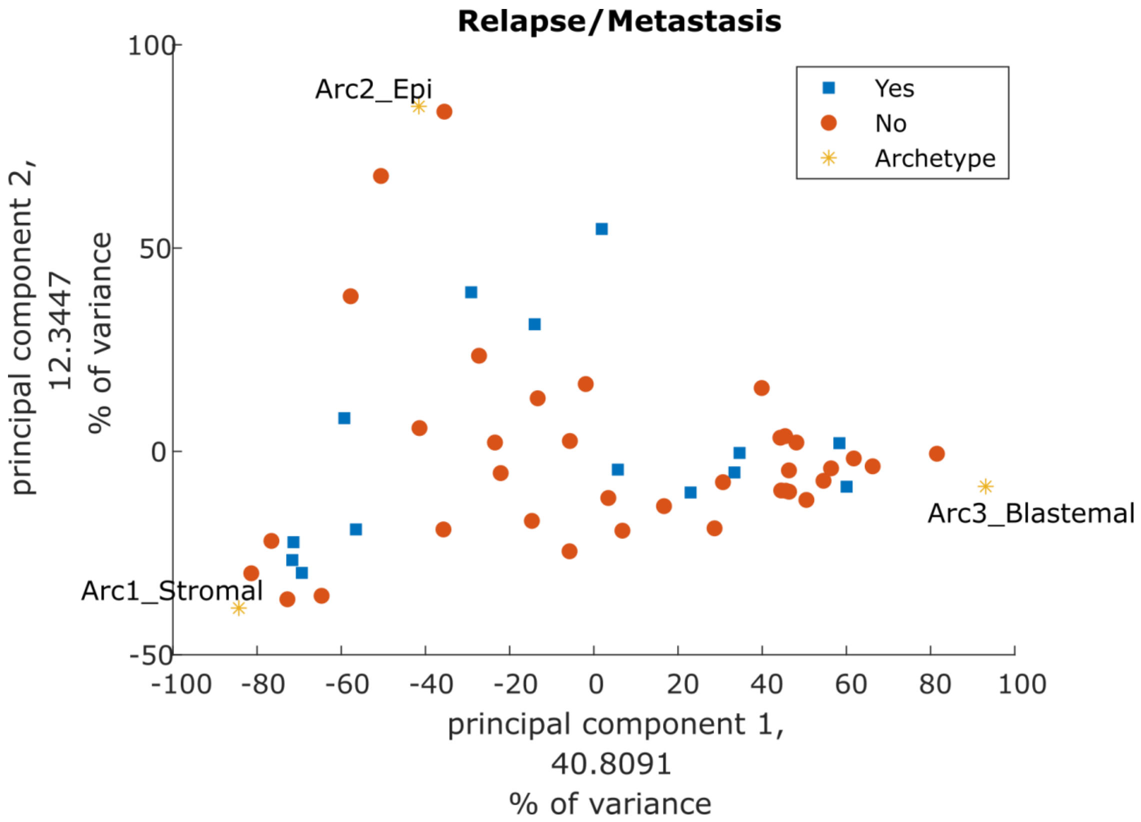


Figure S20: PCA of tumors and archetypes showing the reported status of the tumor (relapse/metastasis).

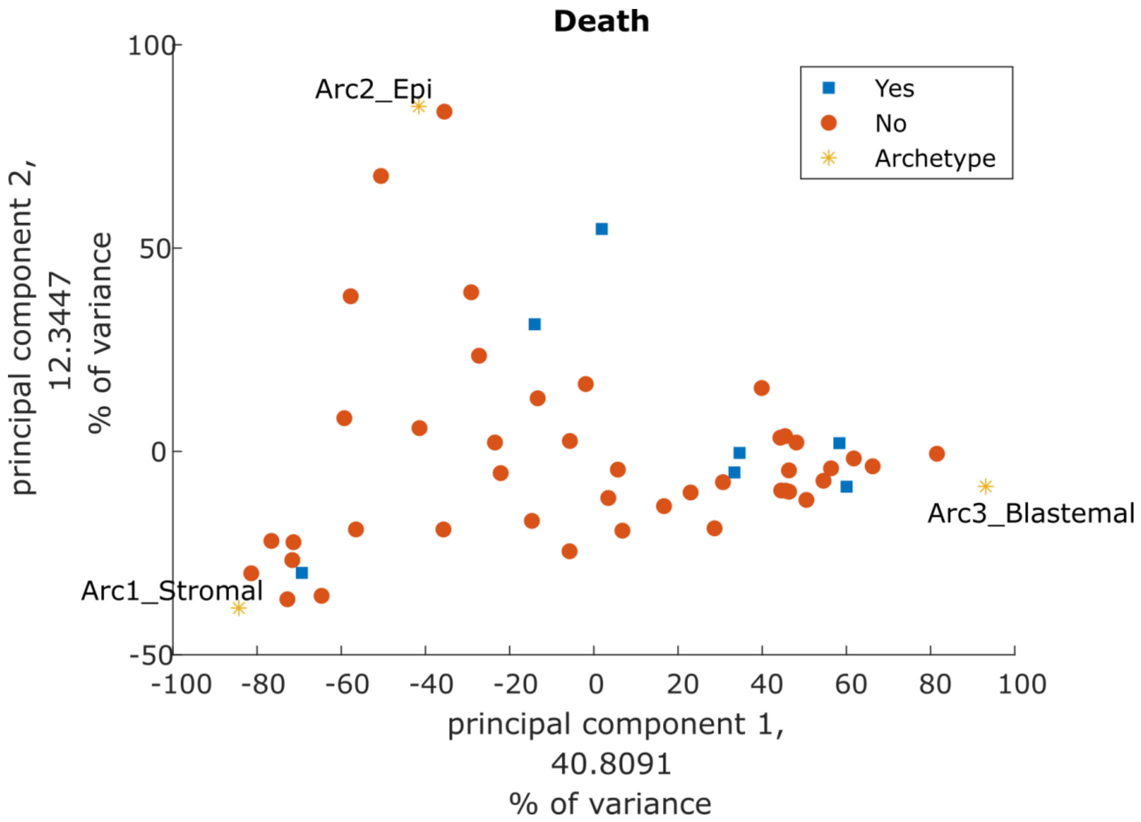


**Figure S21:** PCA of tumors and archetypes showing the reported status of the patient (**death**).

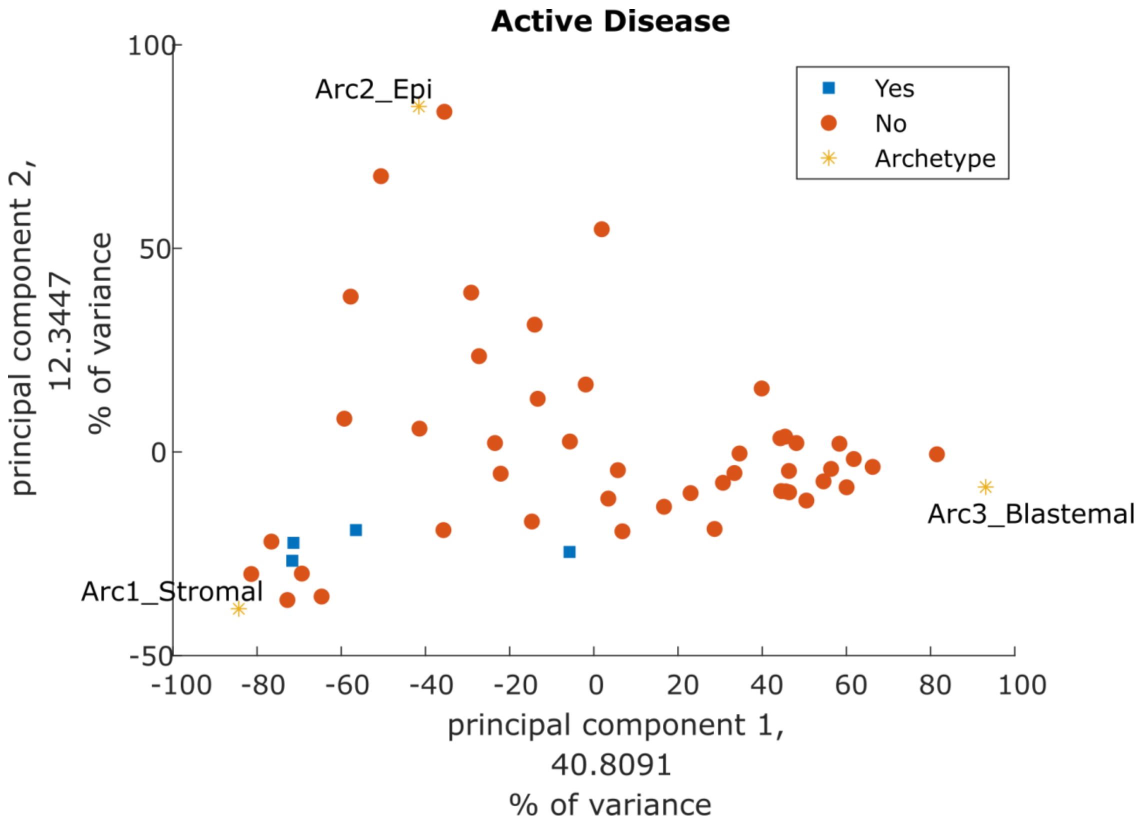


**Figure S22:** PCA of tumors and archetypes showing the reported activity of the tumor (**active disease**). It can be seen that most tumors from patients with active disease in this dataset tend to locate near the stromal archetype.

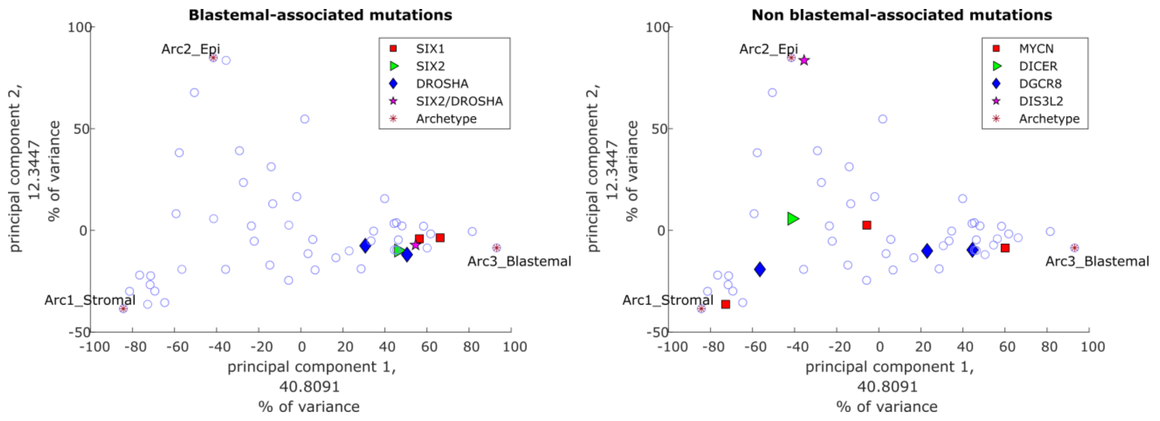


**Figure S23:** PCA of tumors and archetypes showing the reported **mutations**. It can be seen that mutations in the genes SIX1, SIX2, or DROSHA (left panel) tend to cluster in the vicinity of the blastemal archetype. This agrees with the higher incidence of SIX1/2 mutations in tumors with chemotherapy-resistant blastema that was observed by Wegert et al. (Wegert et al., 2015).

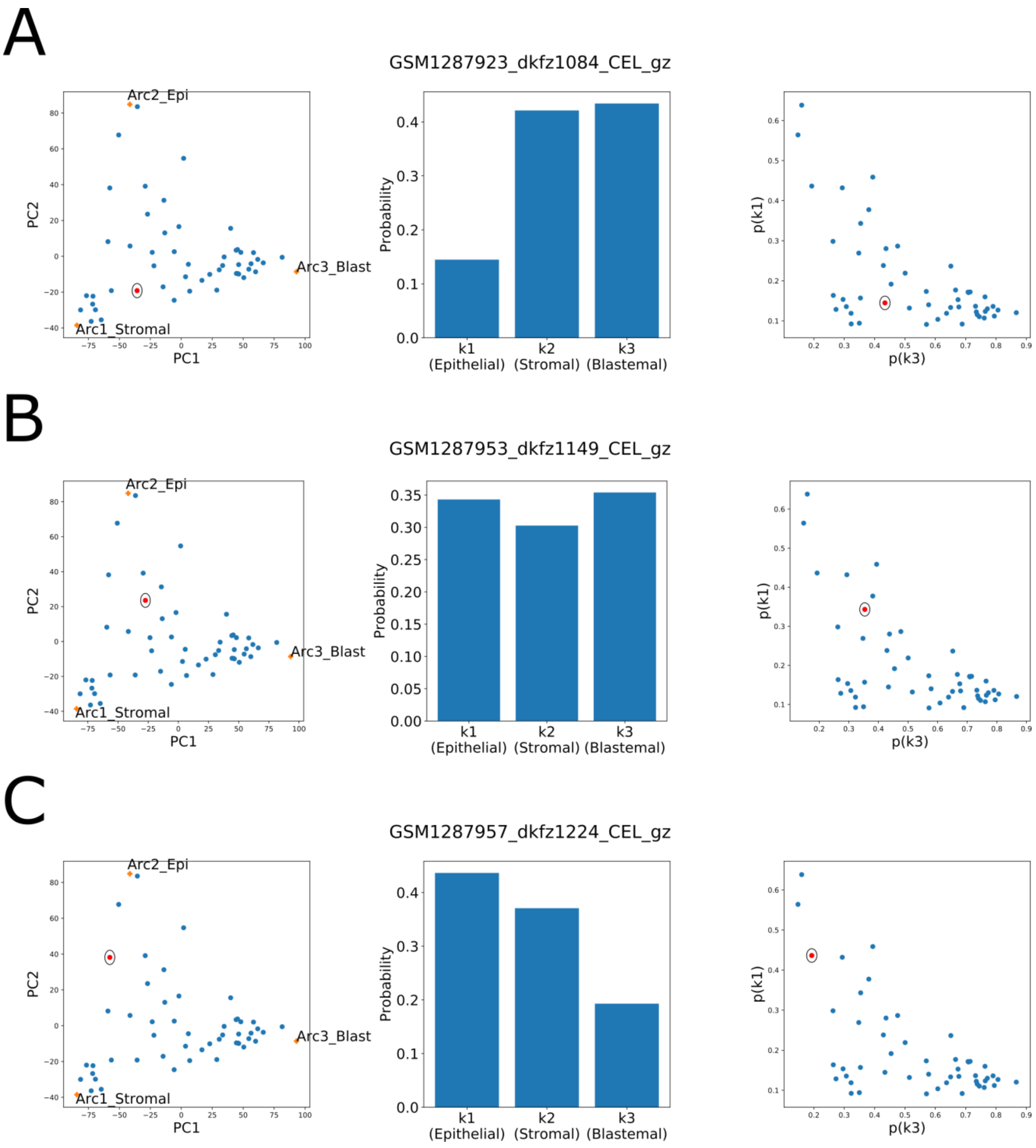


**Figure S24:** LDA topic modeling shows that tumors in-between the vertexes of the triangle-shaped continuum (left panels) are composed of multiple topics (middle panels), in consistency with the topic simplex (right panels).

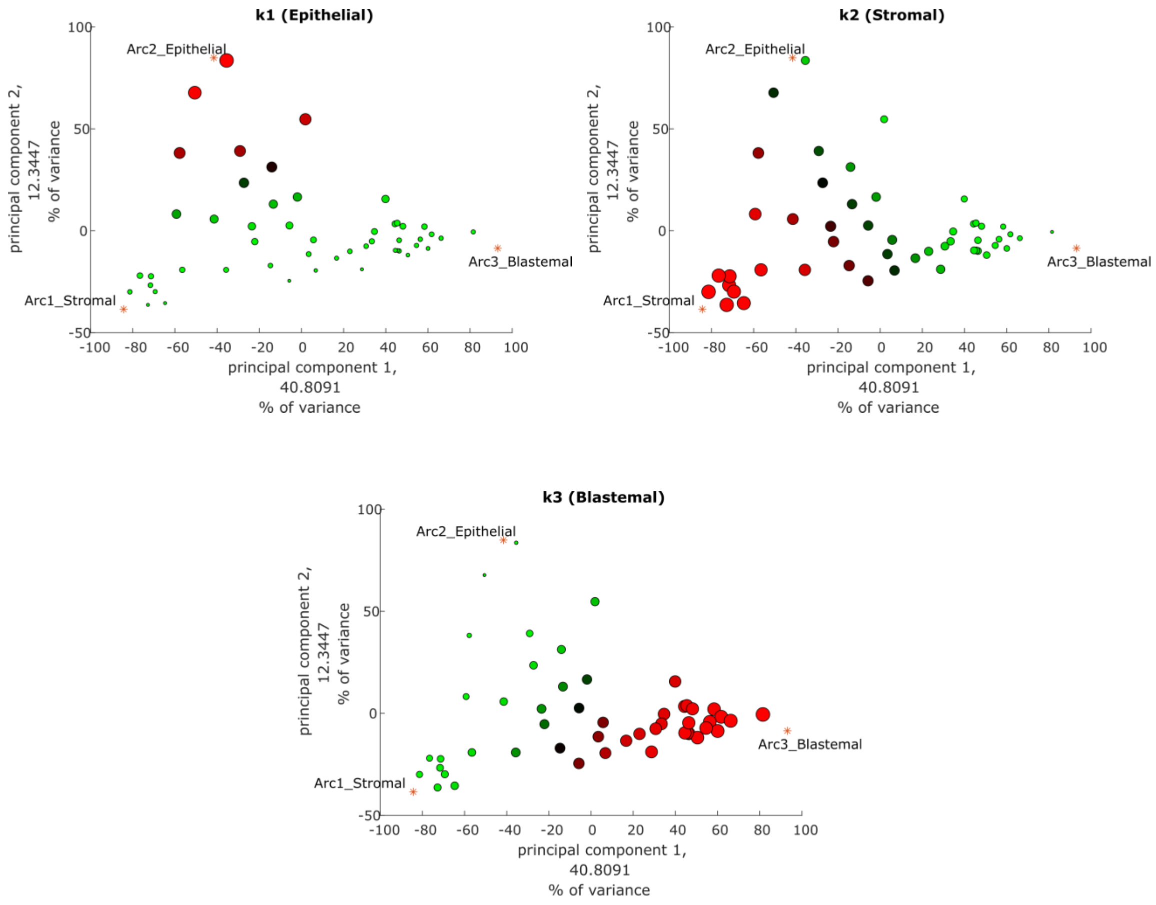


Figure 25: The proportions of each of the three topics are highest towards each of the different vertexes of the triangle-shaped continuum. Shown are PCA plots of the tumors in our dataset. For each topic, large proportion – red/large symbol, small proportion – green/small symbol.

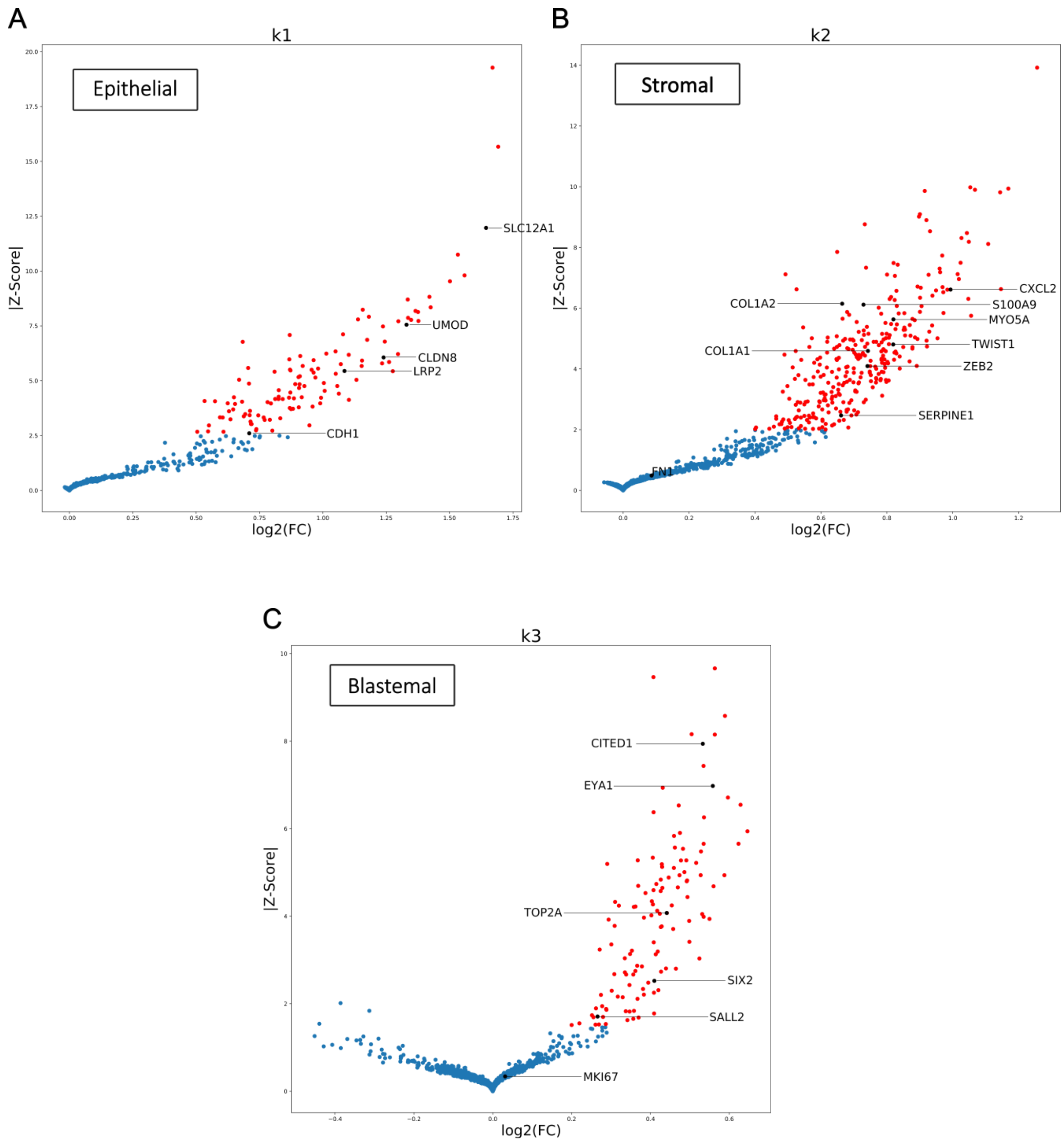


**Figure S26:** The three topics over-express genes related to renal epithelial, stromal, or blastemal characteristics. Shown are volcano plots of the log-fold-change vs Z-score for each of the topics (compared against the null model). We marked genes that are known to be over-expressed in the renal epithelial structures, the un-induced mesenchyme (stroma), and the Cap mesenchyme (blastema) of the fetal kidney.

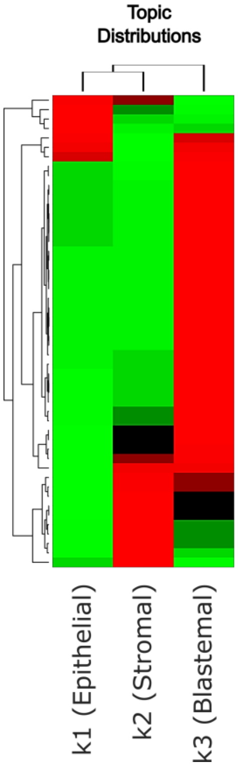


| Sample names | Sex (male/female) | Age (months) | bilateral/Relapse | Histological Type | Histology | Stage | Metastasis at Diagnosis | Relapse/Metastasis | Death | Active Disease | Mutations |
|---|---|---|---|---|---|---|---|---|---|---|---|
| GSM1287957 | Female | 50 | Bilateral | Blastemal | Blastemal | 1 | | | | | |
| GSM1287925 | Male | 58 | | Regressive | Regressive | 1 | | | | | |
| GSM1287948 | Male | 41 | | Blastemal | Blastemal | 2 | y | | | | DIS3L2 c.C2479T:p.Q827X * (het) |
| GSM1287966 | Male | 45 | - | Regressive | Regressive | 1 | | y | | | |
| GSM1287936 | Female | 22 | Bilateral | Blastemal | Blastemal | 3 | y | y | y | | |
| GSM1287954 | Female | 29 | Bilateral | Blastemal | Blastemal | 3 | y | y | y | | |
| GSM1287953 | Female | 82 | Multifocal | Blastemal | Blastemal | 1 | | | | | |
| GSM1287933 | Male | 6 | | Epithelial | Epithelial type | 1 | | | | | |
| GSM1287955 | Female | 153 | - | Blastemal | Blastemal | 1 | | | | | |
| GSM1287958 | Female | 23 | - | Blastemal/xenograft | Blastemal | 3 | | | | | |
| GSM1287924 | Female | NaN | Relapse | Diffuse anaplasia | Diffuse anaplasia | 3 | y | y | y | | |
| GSM1287962 | Female | 20 | - | Blastemal | Blastemal | 1 | | | | | |
| GSM1287959 | Female | 83 | Multifocal | Blastemal | Blastemal | 1 | | | | | |
| GSM1287934 | Female | 148 | | Regressive | Regressive | 1 | | | | | |
| GSM1287960 | Female | 5 | - | Blastemal before chemotherapy | Blastemal | 1 | | | | | |
| GSM1287941 | Male | 34 | Bilateral | Blastemal/focal anaplasia | Blastemal | 1 | | | | | SIX1 c.A530G, p.Q177R (hom) |
| GSM1287944 | Male | 31 | Bilateral | Blastemal | Blastemal | 2 | y | | | | SIX1 c.A530G, p.Q177R (het) |
| GSM1287938 | Male | 52 | - | Diffuse anaplasia | Blastemal | 2 | | y | y | | |
| GSM1287950 | Female | 88 | | Blastemal | Blastemal | 3 | y | y | y | | MYCN c.C130A:p.P44T & c.C131T:p.P44L (hom) |
| GSM1287961 | Female | 55 | | Blastemal | Blastemal | 2 | | | | | SIX2 c.A530G, p.Q177R (het); DROSHA c.G3439A:p.E1147K (het) |
| GSM1287964 | Male | 35 | - | Blastemal | Blastemal | 1 | | | | | |
| GSM1287921 | Female | 35 | | Mixed | Triphasic | 2 | | | | | |
| GSM1287922 | Female | 43 | | Blastemal | Blastemal | 3 | | | | | SIX2 c.A530G, p.Q177R (het) |
| GSM1287943 | Female | 66 | - | Diffuse anaplasia | Blastemal | 3 | | y | y | | |
| GSM1287935 | Female | 29 | | Mixed | Triphasic | 1 | | | | | DROSHA c.G3439A:p.E1147K (het) |
| GSM1287949 | Female | 44 | Bilateral | Blastemal/regressive | Blastemal | 1 | | | | | DGCR8 c.G1552A:p.E518K (hom) |
| GSM1287940 | Male | 8 | - | Blastemal | Blastemal | 1 | | | | | |
| GSM1287970 | Female | 26 | | Blastemal | Blastemal | 2 | | | | | DROSHA c.G3664A:p.E1222K (het) |
| GSM1287945 | Female | 78 | | Blastemal | Blastemal | 1 | | y | | | DGCR8 c.243_244insT:p.K82X (hom) |
| GSM1287963 | Female | 109 | - | Blastemal | Blastemal | 1 | | y | | | |
| GSM1287969 | Female | 85 | - | Blastemal | Blastemal | 3 | | | | | |
| GSM1287952 | Female | 50 | Bilateral | Blastemal | Blastemal | 1 | | | | | MYCN c.C131T:p.P44L (het) |
| GSM1287937 | Male | 12 | - | Blastemal | Blastemal | 1 | | | | | |
| GSM1287942 | Male | 41 | | Blastemal | Blastemal | 1 | | | | | |
| GSM1287951 | Male | 23 | Bilateral | Blastemal | Blastemal | 3 | y | | | | |
| GSM1287956 | Female | 25 | - | Blastemal | Blastemal | 1 | | | | | |
| GSM1287929 | Female | 45 | | Regressive | Regressive | 1 | | | | y | |
| GSM1287928 | Male | 22 | | Mixed | Triphasic | 1 | | | | | |
| GSM1287920 | Female | 41 | | Stromal | Stromal type | 1 | | | | | |
| GSM1287923 | Male | 27 | | Mixed | Triphasic | 1 | | | | | |
| GSM1287946 | Male | 9 | | Blastemal | Blastemal | 1 | | | | | DICER1 c.G5656A:p.G1886R * (het) |
| GSM1287947 | Female | 146 | | Blastemal | Blastemal | 2 | | y | | y | DGCR8 c.G637T:p.E213X (het) |
| GSM1287927 | Male | 37 | | Mixed | Triphasic | 3 | | | | | |
| GSM1287932 | Male | 116 | | Regressive | Regressive | 2 | | | | | MYCN c.C131T:p.P44L (het) |
| GSM1287919 | Male | 124 | | Regressive | Regressive | 3 | y | y | y | | |
| GSM1287918 | Female | 101 | | Necrotic | Completely necrotic | 3 | y | y | | y | |
| GSM1287926 | Female | 4 | | Mixed | Triphasic | 1 | | y | | y | |
| GSM1287930 | Female | 60 | | Regressive | Regressive | - | y | | | | |
| GSM1287931 | Female | 42 | | Necrotic | Completely necrotic | 1 | | | | | |
| GSM1287939 | Male | 82 | - | Regressive | Regressive | 1 | | y | | | |

**Figure S27:** Heatmaps showing the topic distributions of each tumor vs. tumor histological and clinical metadata. It can be seen that tumors with anaplastic histology (diffuse or focal), which is considered least favorable, as well as the single blastemal xenograft in the dataset, and tumors with mutations in the genes SIX1, SIX2, or DROSHA, all contain a large fraction of the blastemal topic.

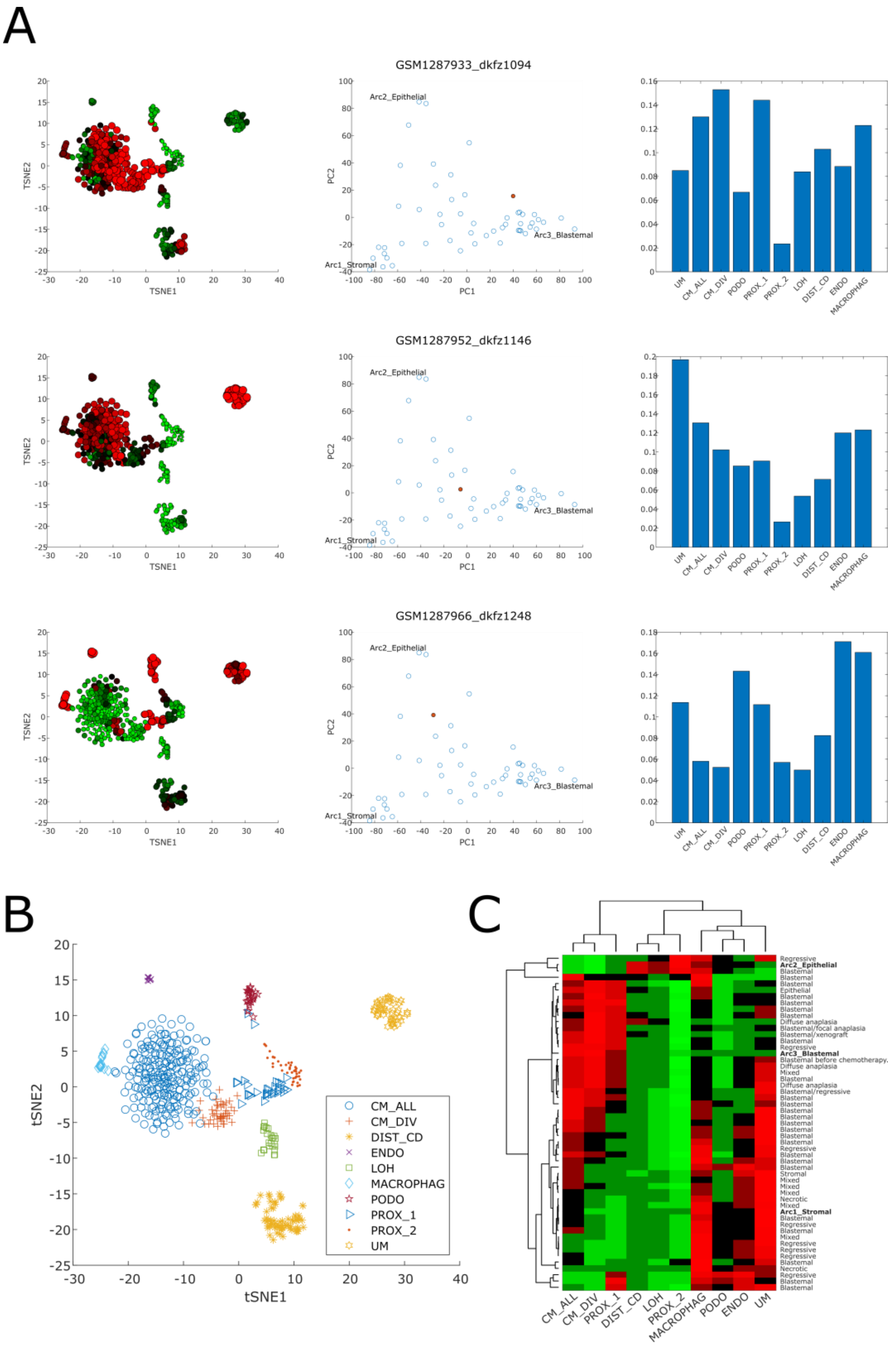


**Figure S28:** Cellular deconvolution indicates that each tumor is composed of a unique mixture of cell populations resembling the un-induced mesenchyme, the cap mesenchyme, or the early epithelial structures of the fetal kidney. (A) Shown is a cellular deconvolution of selected tumors located within the triangle-shaped continuum that is spanned by the stromal, blastemal, and epithelial archetypes. It can be seen that tumors located in-between the three archetypes are composed from a mixture of heterogeneous cell types, for example, cells from the Cap mesenchyme and the renal epithelium (top panel), cells from the Cap mesenchyme and the un-induced mesenchyme (middle panel), and cells from the un-induced mesenchyme and the renal epithelium (bottom panel). This is in contrast to the archetypes that are each predominantly composed of a single cell type (the un-induced mesenchyme, the Cap-mesenchyme, or epithelial cells, see Fig. 4). (B) The different cell populations marked on a tSNE plot of the reference single cell RNAseq dataset from the developing mouse fetal kidney that was used for cellular deconvolution (CM – Cap mesenchyme, DIST_CD – distal tubule and collecting duct, ENDO – endothelial, LOH – Loop of Henle, MACROPHAG – macrophages, PODO – podocytes, PROX_1 – early epithelial structures such as C/S-shaped bodies, PROX_2 – proximal tubule, UM – un-induced mesenchyme). (C) A heatmap of the cell type proportions from which each tumor is composed, as predicted by cellular deconvolution.

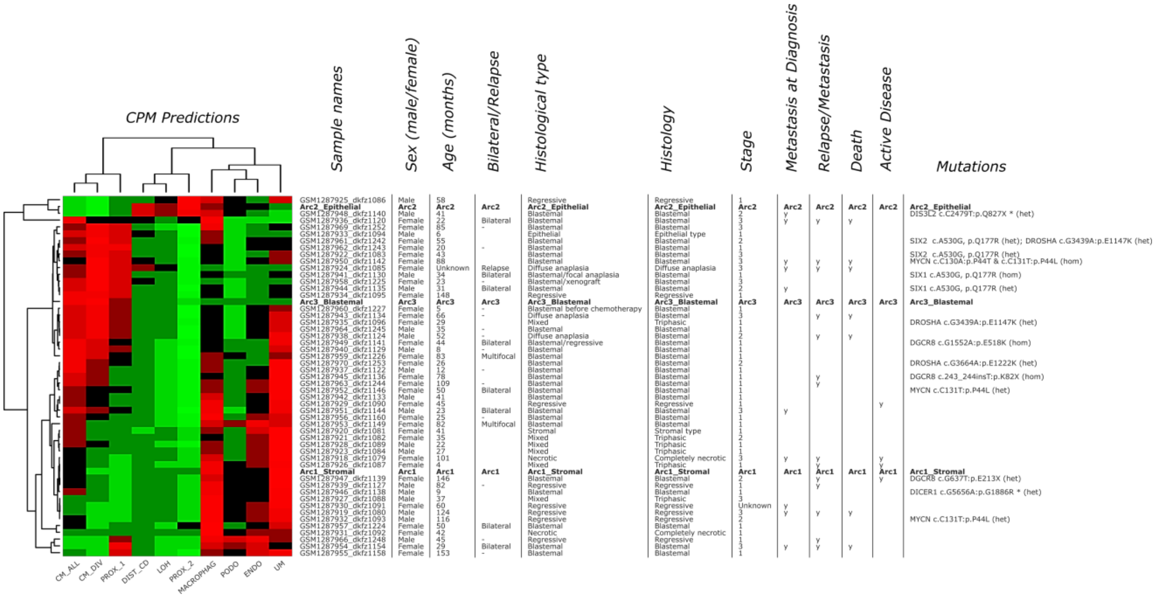


**Figure S29:** A heatmap of the cell type proportions from which each tumor is composed, as predicted by cellular deconvolution, along with tumor histological and clinical metadata. It can be seen that most of the tumors with reported blastemal histology contain a significant proportion of cells resembling those of the Cap mesenchyme (CM_ALL), as expected. Likewise, it can be seen that tumors with anaplastic histology (diffuse or focal), as well as the single blastemal xenograft in our dataset, and also tumors reported to contain mutations in the genes SIX1, SIX2, or DROSHA, contain a significant fraction of cells resembling the cycling Cap mesenchyme cells (CM_DIV) in the fetal kidney. This is in agreement with the findings of Wegert *et al*. [2] that blastemal type Wilms tumors with mutations in SIX1 or SIX2 have a have a gene expression signature of proliferation and kidney progenitors. On the other hand, it can be seen that tumors reported to have triphasic/mixed or regressive histology contain high proportions of cells resembling the un-induced mesenchyme (UM).
